# WhaleSonic: Recordings of cetacean vocal behaviour with hydrophone arrays in Northern Norway

**DOI:** 10.64898/2026.11.02.763576

**Authors:** Jörg Rychen, Arvid Berg, Jerome Cohen, Isabel Driscoll, Alexander Eckerle, Valeria Falletta, Olga Filatova, Lena Gies, Pascale Hatt, Adrian Hiss, Franziska Krimmer, Corinna Lorenz, Joseph G. Mine, Charlotte Mommsen, Florian S. Schmidt, Rik L.E.M. Ubaghs, Kaja Wierucka, Richard Hahnloser

## Abstract

Killer whales, humpback whales, fin whales, and sperm whales converged in a fjord system in Northern Norway during the 2023–2024 winter, where whale watching and fishing vessels add to the acoustic scene. Cetaceans are intensely studied for their acoustic behaviour and communication, and the co-occurrence of four species in confined waters offers a unique opportunity to record them with hydrophone arrays. We present WhaleSonic, a raw acoustic dataset recorded with two synchronised array systems. Four drifting buoys, each carrying a tetrahedral hydrophone array, captured 78 hours of calibrated recordings; seven bottom-mounted hydrophones yielded 52 days of continuous data. Both systems are synchronised to UTC (***<***1 μs and 3–7 ms) and positioned to 2 m accuracy. Transmission experiments and sound velocity measurements provide in-situ ground truth. All data are in physical units and a unified coordinate system, self-documented in HDF5 files, and published with open-source tools. WhaleSonic supports the development of multichannel algorithms for source separation, localisation, sound field analysis, and group communication in free-ranging cetaceans.

## Background & Summary

The evolutionary origin of human language remains one of the most fundamental open questions in science. Studying vocal communication in other species is one of the few available approaches to understanding what drives the evolution of communication systems [1]. Linguistic concepts such as combinatoriality, compositionality, and sequential structure can be applied comparatively across animal vocal systems, revealing shared building blocks whose phylogeny can be traced [2]. Cetaceans are a particularly compelling model: they rely on sound as their primary sense, have evolved large brains and complex social structures, and are capable of vocal production learning [3]. Machine learning is now being applied to analyse animal communication at scale [4, 5], and multichannel acoustic datasets can provide important advantages for these efforts, as we lay out below. We first introduce the four cetacean species recorded in this dataset, each with diverse acoustic behaviours.

Killer whales (*Orcinus orca*) produce group-specific repertoires of stereotyped calls that form vocal dialects, diverging between matrilineal families and showing geographic variation across North Atlantic populations that track herring migration routes [6–9]. Norwegian killer whales cooperatively herd herring (*Clupea harengus*) into tight balls near the surface, a behaviour known as carousel feeding [10], using tail slaps to debilitate prey [11]. Analysis of Norwegian killer whale calls suggests they are composed of shared segments that can be recombined, hinting at combinatorial structure [12]. Using a towed hydrophone array to attribute calls to individuals, Miller et al. [13] discovered call-type matching in vocal exchanges between group members.

Humpback whales (*Megaptera novaeangliae*) are among the most acoustically elaborate cetaceans. Males produce long, hierarchically structured songs whose patterns change progressively within and between seasons [14], with evidence of underlying syntactic structure [15]. Singing in humpbacks was traditionally associated with tropical breeding grounds, but has recently been documented on subarctic feeding grounds in Norway [16], where humpback whales aggregate alongside killer whales to forage on overwintering Norwegian spring-spawning herring [17].

Fin whales (*Balaenoptera physalus*) produce stereotyped low-frequency songs consisting of 20 Hz pulses, often paired with a higher-frequency component near 130 Hz, repeated at metronomic intervals of 10 s to 40 s for hours [18]. Only males are known to sing [19]. Pulse patterns and inter-note intervals vary between ocean basins and shift over years [20], providing acoustic markers of population identity.

Sperm whales (*Physeter macrocephalus*) produce powerful broadband clicks for echolocation during deep foraging dives, and communicate using rhythmically patterned click sequences known as codas [21]. Their clicks are highly directional, and their three-dimensional beam pattern has been characterised using hydrophone arrays [22]. Male sperm whales forage year-round in Northern Norway [23].

These four species converged in the Kvænangen fjord system of Northern Norway during the 2023–2024 winter, alongside vast aggregations of herring overwintering in the fjords for two to three months [17, 24, 25]. This brief seasonal convergence creates an exceptional setting: species with fundamentally different acoustic behaviours co-occur in confined waters and can be recorded simultaneously. The narrow, steep-walled fjords also create a complex acoustic environment with strong reverberation, where vocalisations of multiple species overlap in time and space.

Studying vocal communication within a group requires separating the signal of each individual. In the literature, two main approaches have been pursued. Acoustic recording tags (DTags) attached to animals are a standard tool in cetacean research, delivering rich multi-sensor data [26]. When two sperm whales were tagged simultaneously, their coda exchanges could be analysed as conversations for the first time [27]; similarly, tagged humpback whale mother-calf pairs revealed acoustic exchanges during migration [28]. For small animals such as songbirds, tagging entire groups is feasible and has enabled the analysis of communication networks in groups of up to eight individuals [29]. For cetaceans, however, tagging all or even most animals in a group remains impractical. The second approach—and the one we use here—is to deploy hydrophone arrays, which require no animal contact: they record all animals simultaneously, making them well suited for studying group communication while minimising disturbance. The trade-off is that calls can only be attributed to different individuals based on spatial separation—a signal processing challenge that this dataset can address.

Most bioacoustic analysis tools—detection, classification, species identification— operate on single-channel audio. Even when multichannel recordings are available, detection and classification are typically performed per channel, with spatial information exploited only downstream for localisation [30]. Neural network approaches to source separation in bioacoustics exist but likewise operate on single channels [31]. Yet when multiple animals vocalise simultaneously, spectral cues alone are often insufficient to disentangle overlapping calls, which is where temporal cues from multi-channel recordings can make a difference.

With multiple spatially distributed hydrophones, the same vocalisation arrives with different time delays at each receiver, providing spatial information for source separation and localisation. Hydrophone arrays in cetacean research span a wide range of configurations: towed arrays [30, 32], drifting surface buoys [33, 34], deep-water vertical arrays [35], cabled coastal arrays [36, 37], bottom-mounted arrays [38, 39], and compact tetrahedral arrays [40, 41]. PAMGuard [42] provides comprehensive array processing tools for a wide range of these geometries. The primary use of array recordings has been to localise and track animals—yielding dive profiles, movement patterns, and habitat use. Recent frameworks for automated multi-animal tracking [43, 44] demonstrate the growing maturity of these methods.

Another use of hydrophone arrays is emission beam characterisation: measuring how animals radiate sounds directionally. Au et al. [45] first characterised the transmitting beam of captive dolphins. In the field, Miller [46, 47] measured the mixed-directionality of killer whale stereotyped calls using a towed array, sperm whale click beam patterns were resolved with bottom-mounted arrays [22, 48], and purpose-built vertical arrays have been deployed for beam measurements of deep-diving odontocetes in the wild [49, 50].

Arrays have also been used to study vocal communication directly. Miller et al. [13] attributed calls to individual killer whales from a towed array and discovered call-type matching in vocal exchanges. Poupard et al. [51] used a compact shore-based tetrahedral array to estimate intra-group call rates. Tenorio-Hallé et al. [52] distinguished concurrent humpback whale singers using direction-of-arrival information from vector sensors. These studies demonstrate that spatial information from arrays can advance the analysis of vocal behaviour beyond what single-channel methods can achieve. WhaleSonic was recorded specifically with the analysis of cetacean group communication in mind.

This dataset was designed as the interface between observation and analysis. On the observation side we managed recording technology, field operations, and data management; the consolidation pipeline resolves all instrument-specific complications and delivers measurements in physical units. On the analysis side are the research questions, addressed with tools such as detection, classification, localisation, and source separation. This paper describes only the observation. Measurements do not change; analysis built upon them will evolve as algorithms improve. The aim was to depict the acoustic environment accurately, providing a shared empirical foundation for different research groups to apply and compare their own methods.

The WhaleSonic dataset was recorded in winter 2023–2024 near Skjervøy, Northern Norway (Fig. 1). We maximised effort on two dominant error sources in array processing: clock offsets and receiver positions [53]. Four drifting recorder buoys, each carrying a tetrahedral hydrophone array, captured 82 recordings totalling 78 hours with sub-microsecond synchronisation to UTC and 1.9 m positioning accuracy. Seven bottom-mounted hydrophones yielded 52 days of continuous data with 2.5 ms to 6.6 ms RMS timing and ~2 m acoustically determined positions. Sound emission tests and 118 sound velocity profiles provide in-situ ground truth for transmission and propagation studies [54]. The full timeline is shown in Fig. 2.

**Fig. 1.**
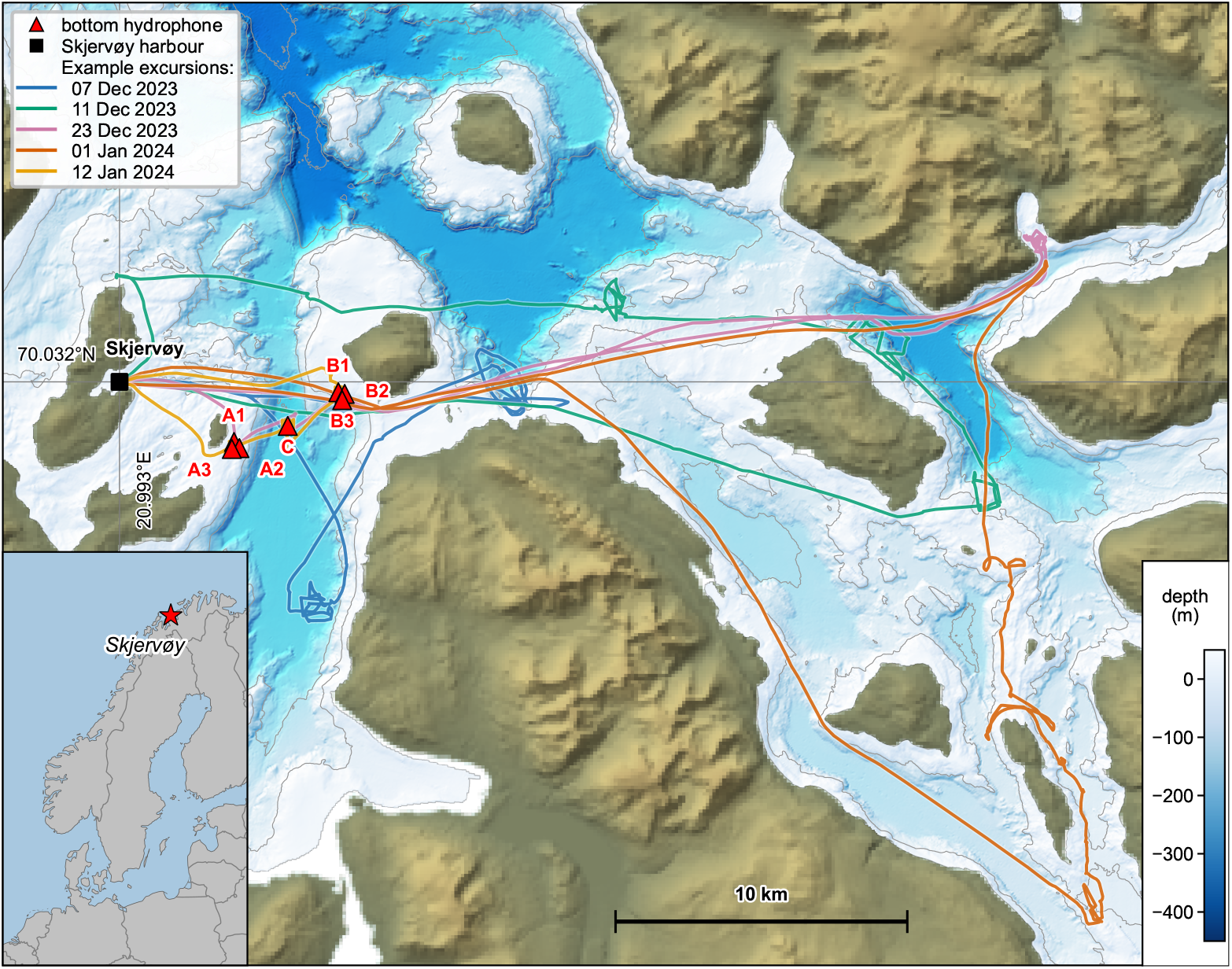
Study area and recording coverage. Kvænangen fjord system near Skjervøy, Northern Norway, with bathymetry (colour scale). Red triangles mark the seven bottom-mounted hydrophones (clusters A1–A3, B1–B3, and isolated position C). Coloured lines show research vessel excursion tracks for five example recording days, illustrating the spatial extent of buoy deployments across the study area. The black square marks Skjervøy harbour. Inset: location of the study area within Norway.

**Fig. 2.**
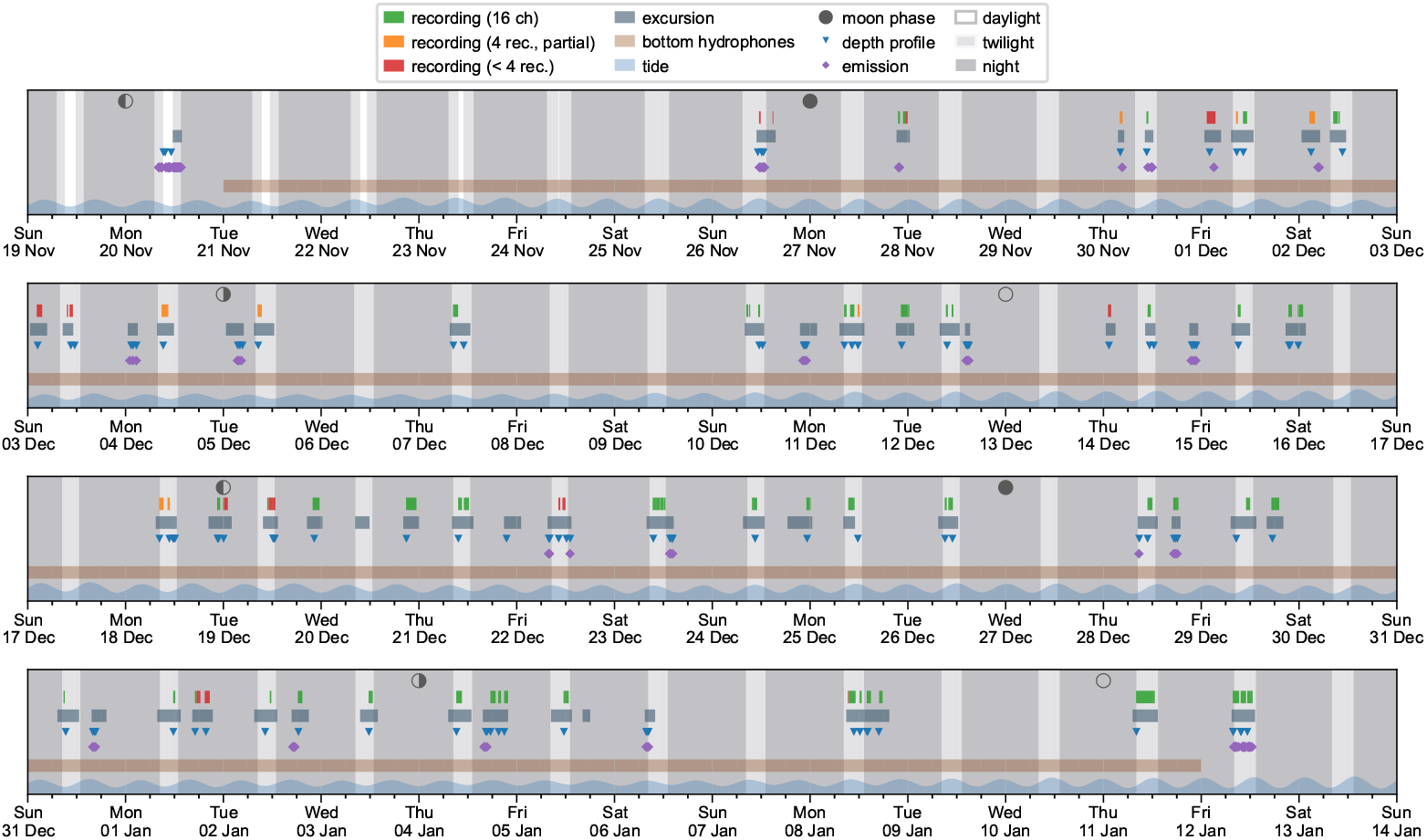
Expedition timeline. Multi-row calendar view of all dataset components and expedition activities across the eight-week field season (19 November 2023 to 13 January 2024). Each row spans two weeks. From top to bottom within each row: buoy recordings (coloured by array completeness: green = all 16 channels, orange = 4 recorders with partial channels, red = fewer than 4 recorders), research vessel excursions, sound velocity profiles, underwater speaker emissions, and continuous bottom hydrophone coverage. Tide level is shown along the bottom edge. Background shading indicates the polar night cycle at 70°N (unshaded = daylight, light = civil twilight, dark = night), and moon phase symbols mark cardinal phases along the top.

The dataset is published following FAIR principles [55, 56], accompanied by the open-source `whalesonic` Python package for data access, visualisation, and export. All recordings are stored in self-describing containers (HDF5 and FLAC with YAML meta-data) that embed array geometry, calibration, and sound velocity data. We publish these raw recordings to foster the development of multichannel bioacoustic methods, and hope that subsequent analyses will likewise be shared openly for the benefit of the field.

A companion dataset from a follow-up deployment in the 2025–2026 season (Tverrfjord 2025/2026) [57] provides an additional 53 days of continuous seven-channel bottom hydrophone recordings from the same fjord system, recorded with the same instruments.

## Methods

### Recording area

All recordings were made in the waters around Skjervøy, a fishing village in the Kvæ-nangen fjord system of Northern Norway at ~70°N, ~21°E (Fig. 1). At this latitude, the sun remains below the horizon from late November to mid-January (56 days of polar night), with only about four hours of civil twilight at the winter solstice—when the sun’s max elevation is −2.5°. The full moon, conversely, can remain continuously above the horizon for several days around each full moon phase, providing significant illumination during the polar night. Despite the high latitude, the North Atlantic Currrent keeps the fjords ice-free with winter sea surface temperatures of 4 °C to 6 °C, warmer than at comparable latitudes in the western Atlantic or Siberian Arctic. Norwegian spring-spawning herring shifted their overwintering area to the Kvænangen region around 2017 [24], and the seasonal presence of cetaceans has since turned the area into a high-traffic zone: a diverse whale-watching industry [58] operates alongside industrial purse-seine fisheries [59] and scientific expeditions each winter.

### Environmental data and coordinate system

All times in the dataset are expressed in UTC. All spatial coordinates are expressed in a local stereographic projection centred on the entrance of Skjervøy harbour (70.032°N, 20.993°E; Fig. 1) that converts WGS84 coordinates to Cartesian coordinates in metres. The projection is defined in PROJ [60] format in the dataset’s HDF5 files and in the `whalesonic` Python package.

Seafloor depth data were obtained from the Kartverket DTM50 digital terrain model [61] (50 m resolution) and interpolated to a uniform 10 m grid in the local coordinate system. Tidal water levels were obtained from the Kartverket water level API [62].

### Expedition operations

The bottom hydrophones were deployed on 20 November 2023 and recovered on 12 January 2024, framing a 53-day campaign (see Bottom hydrophone array below). Throughout this period, excursions were conducted from a base in Skjervøy using a rigid-hull motorboat (“Malin 5”, 7.5 m length, 150 hp outboard engine). The aim of each excursion was to deploy the four drifting buoy recorders around a scene with promising acoustic activity. Two teams operated in alternation, each consisting of an experienced skipper and two to three field scientists. One team covered the daylight and twilight hours, the other team went out in darkness. The day team could visually search for animals; the night team approached predetermined spots with presumed activity or used occasional underwater listening to find a suitable deployment location. It was up to each team to decide where and when to deploy the buoys; there was no prescriptive protocol for effort allocation, other than to record a set of soundscapes as diverse as possible. While the buoys were recording, the day team could fly drones and conduct photo-identification. After each excursion, all instruments were brought back to a shore base for maintenance, battery recharging, and data download. The expedition was designed for high throughput, building on experience from similar deployments since 2015. Over the course of the expedition, 61 excursions were completed.

### Buoy recorder system

The buoy recording system consists of four free-drifting surface buoys (Rec-A through Rec-D), each with a tetrahedral hydrophone array suspended below in the water. The design builds on the large-aperture array concept of Møhl et al. [33] and Miller & Dawson [63], extending it with nested tetrahedral sub-arrays that enable both interbuoy localisation over hundreds of metres and direction-of-arrival estimation from each tetrahedron (Table 1).

**Table 1.**
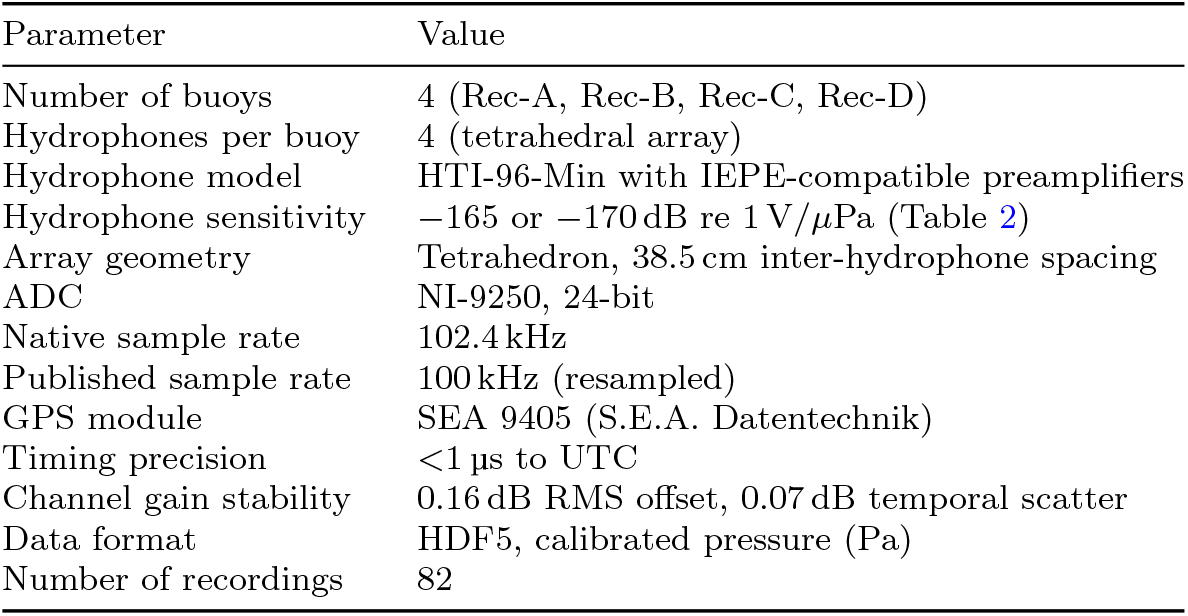
Buoy recorder specifications.

| Parameter | Value |
|---|---|
| Number of buoys | 4 (Rec-A, Rec-B, Rec-C, Rec-D) |
| Hydrophones per buoy | 4 (tetrahedral array) |
| Hydrophone model | HTI-96-Min with IEPE-compatible preamplifiers |
| Hydrophone sensitivity | −165 or −170 dB re 1 V/*µ*Pa (Table 2) |
| Array geometry | Tetrahedron, 38.5 cm inter-hydrophone spacing |
| ADC | NI-9250, 24-bit |
| Native sample rate | 102.4 kHz |
| Published sample rate | 100 kHz (resampled) |
| GPS module | SEA 9405 (S.E.A. Datentechnik) |
| Timing precision | <1 µs to UTC |
| Channel gain stability | 0.16 dB RMS offset, 0.07 dB temporal scatter |
| Data format | HDF5, calibrated pressure (Pa) |
| Number of recordings | 82 |

The buoys were purpose-built and iteratively optimised for durability and rapid deployment. All electronics are housed in a cylindrical tube (20 cm diameter, 80 cm height) made from high-density polyethylene (PE-HD), a material that is strong yet shock-resistant even at low temperatures. At ~10 kg, each recorder can be carried by a single person and deployed from a small motorboat. Surface wave motion is the primary source of mechanical noise on a drifting buoy; a suspension consisting of a bungee cord, lead weights, and a hydrodynamic damping disk decouples this motion from the hydrophone array (Fig. 3a).

**Fig. 3.**
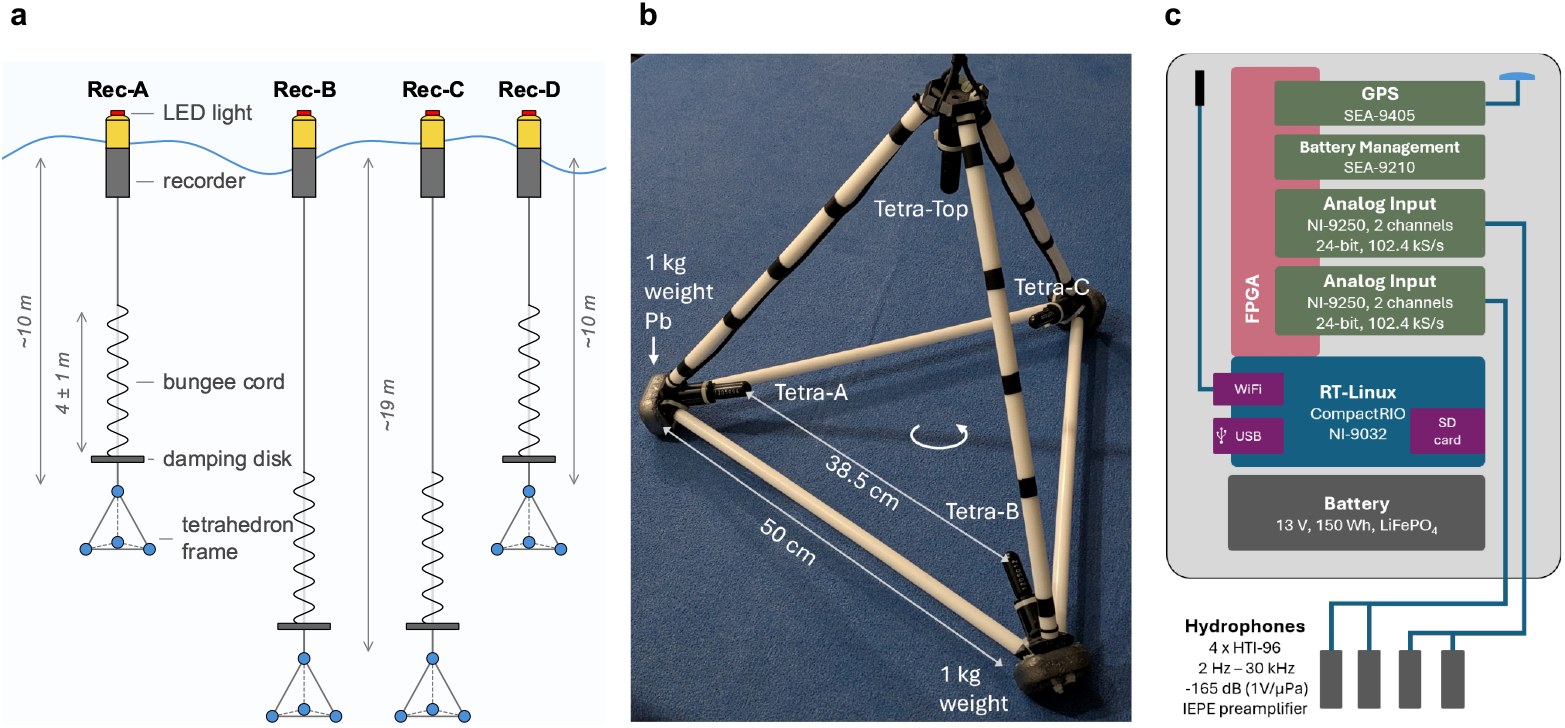
Drifting buoy recorder system. **a** Schematic of the four drifting recorder buoys (Rec-A through Rec-D) with two different tetrahedron depths. Each tetrahedron hangs below the buoy, connected via a bungee cord (4 m ± 1 m) and a damping disk to decouple wave-induced buoy motion and keep the hydrophone array quiet. **b** Tetrahedral hydrophone frame with four HTI-96 hydrophones at positions Tetra-Top, Tetra-A, Tetra-B, and Tetra-C, and three lead ballast weights. The measured inter-hydrophone spacing is 38.5 cm. **c** Block diagram of the recorder electronics. Two dual-channel analogue input modules (NI-9250), a GPS module (SEA-9405), and a battery management module (SEA-9210) are connected to the FPGA of a CompactRIO controller (NI-9032) running RT-Linux. The system is powered by a 13 V 150 Wh LiFePO_4_ battery.

During each excursion, the four buoys were deployed in an approximate square formation with side lengths of 100 m to 800 m. During recording, the boat engine was switched off and the vessel drifted with the wind. The buoys drifted freely for recordings lasting from a few minutes up to 4.5 h. A waterproof LED warning light on each buoy enabled retrieval in the dark during the polar night. Operating limits were ~2 m wave height and ~30 km*/*h wind speed.

#### Tetrahedral hydrophone array

Each buoy carried a tetrahedral array of four hydrophones mounted at the corners of the tetrahedron, pointing inward toward the centre (Fig. 3b). The frame was constructed entirely from plastic (acetal rods, 20 mm diameter, 50 cm edge length) to be nearly acoustically transparent in water. The measured inter-hydrophone spacing was 38.5 cm. The inward-pointing geometry protects the hydrophones from direct collisions and preserves the tetrahedral symmetry of the array response.

A 1 kg lead weight was attached to each of the three lower corners, and the lifting point was carefully adjusted so that the tetrahedron hung level (*<*1° from horizontal). The last 5 m of cable were looped and attached to a straight bungee cord that was 3 m unloaded, 4 m under the weight of the submerged tetrahedron, extensible to 5 m. ~50 cm above the tetrahedron, a horizontal damping disk (20 cm diameter, 1 cm thick, with a grid of drilled holes) served as a hydrodynamic damper to suppress resonances in the suspension system. This critically damped suspension kept the tetrahedron stable and free of mechanical knocks in up to 2 m waves. The azimuthal orientation (heading) of the tetrahedron is not measured and must be inferred from the acoustic signals.

The array was suspended below the buoy at depths that depended on the cable configuration: Rec-A and Rec-D at ~10 m, Rec-B and Rec-C at ~19 m. During deployment, the two deeper buoys (B, C) were placed on one diagonal of the square formation and the two shallower ones (A, D) on the other, so that the four tetrahedrons together formed a large-scale tetrahedral geometry for three-dimensional localisation.

#### Recording electronics

The hydrophones (HTI-96-Min, High Tech Inc.; usable bandwidth 2 Hz–30 kHz) were connected via IEPE-compatible preamplifiers to the recording electronics. Each hydrophone was supplied with a factory calibration certificate specifying its individual sensitivity; Table 2 lists all 16 channels with their serial numbers and calibrated sensitivities. The two deeper tetrahedrons (Rec-B, Rec-C at ~19 m) have a nominal sensitivity of −165 dB re 1 V*/*µPa, while the two shallower ones (Rec-A, Rec-D at ~10 m) of −170 dB. The recordings are published as sound pressure in Pa, converted using the individual factory sensitivities.

**Table 2.**
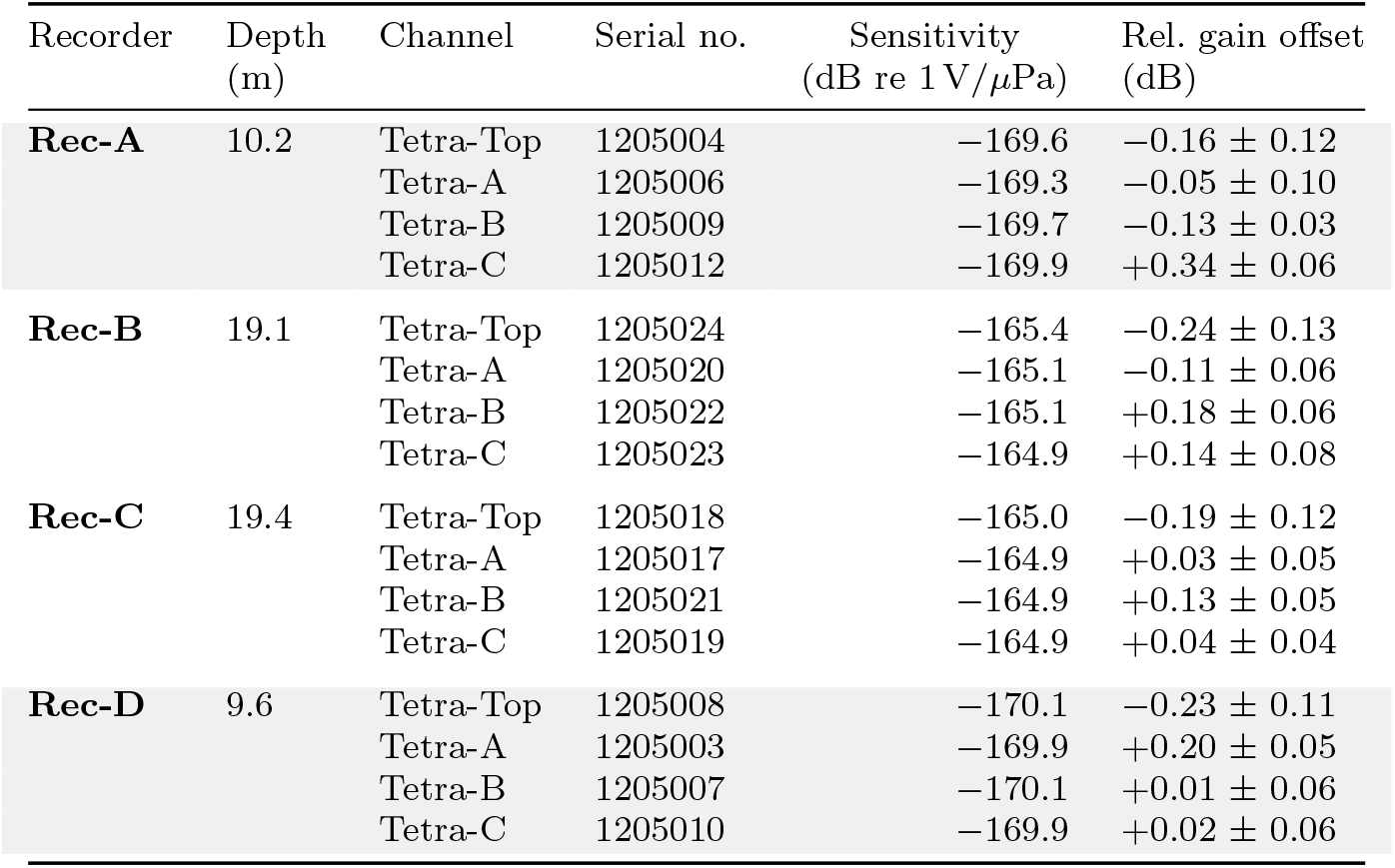
Buoy hydrophone inventory. Individual factory-calibrated sensitivities and measured relative gain offsets for all 16 channels, grouped by recorder. Relative gain offsets are mean ± standard deviation relative to the geometric mean of each tetrahedron, measured across *N* = 63–79 recordings per recorder (see Channel gain stability). Shaded rows indicate the shallow tetrahedrons (~10 m); unshaded rows the deep tetrahedrons (~19 m).

| Recorder | Depth<br>(m) | Channel | Serial no. | Sensitivity<br>(dB re 1 V/*μ*Pa) | Rel. gain offset<br>(dB) |
|---|---|---|---|---|---|
| **Rec-A** | 10.2 | Tetra-Top | 1205004 | −169.6 | −0.16 ± 0.12 |
|  |  | Tetra-A | 1205006 | −169.3 | −0.05 ± 0.10 |
|  |  | Tetra-B | 1205009 | −169.7 | −0.13 ± 0.03 |
|  |  | Tetra-C | 1205012 | −169.9 | +0.34 ± 0.06 |
| **Rec-B** | 19.1 | Tetra-Top | 1205024 | −165.4 | −0.24 ± 0.13 |
|  |  | Tetra-A | 1205020 | −165.1 | −0.11 ± 0.06 |
|  |  | Tetra-B | 1205022 | −165.1 | +0.18 ± 0.06 |
|  |  | Tetra-C | 1205023 | −164.9 | +0.14 ± 0.08 |
| **Rec-C** | 19.4 | Tetra-Top | 1205018 | −165.0 | −0.19 ± 0.12 |
|  |  | Tetra-A | 1205017 | −164.9 | +0.03 ± 0.05 |
|  |  | Tetra-B | 1205021 | −164.9 | +0.13 ± 0.05 |
|  |  | Tetra-C | 1205019 | −164.9 | +0.04 ± 0.04 |
| **Rec-D** | 9.6 | Tetra-Top | 1205008 | −170.1 | −0.23 ± 0.11 |
|  |  | Tetra-A | 1205003 | −169.9 | +0.20 ± 0.05 |
|  |  | Tetra-B | 1205007 | −170.1 | +0.01 ± 0.06 |
|  |  | Tetra-C | 1205010 | −169.9 | +0.02 ± 0.06 |

The recording electronics (Fig. 3c) were built around a CompactRIO controller (NI-9032, National Instruments) with four slots for standardised input/output modules: two NI-9250 two-channel analogue input modules (24-bit, instrument-grade with calibrated voltage reading) sampling the four hydrophone channels at a native sample rate of 102.4 kHz; a GPS module (SEA-9405, S.E.A. Datentechnik) providing time and position at 4 Hz; and a battery management module (SEA-9210) monitoring the battery and a leak detector (BlueRobotics). All modules were connected to the controller’s on-board Field-Programmable Gate Array (FPGA) for low-latency data streaming in reconfigurable hardware logic. The controller ran a real-time Linux (RT-Linux) operating system that streamed data to an SD card and provided a web interface, allowing instrument status display and control from a mobile phone via WiFi. The FPGA logic, the real-time controller application, and the web interface were all developed in Lab-VIEW (National Instruments), an integrated system design platform. The system was powered by a 13 V 150 Wh LiFePO_4_ battery, sufficient for more than eight hours of continuous operation.

#### Synchronisation and resampling

A replica of GPS time running on the FPGA (Timekeeper, National Instruments) was continuously disciplined at 4 Hz to within ±100 ns of the GPS reference (Fig. 4a). The Timekeeper provides a GPS-derived timestamp at every clock cycle of the FPGA (40 MHz, 25 ns), enabling a per-sample GPS timestamp for each ADC sample. Since each buoy carries an independent GPS receiver, inter-buoy synchronisation is better than 1 µs.

**Fig. 4.**
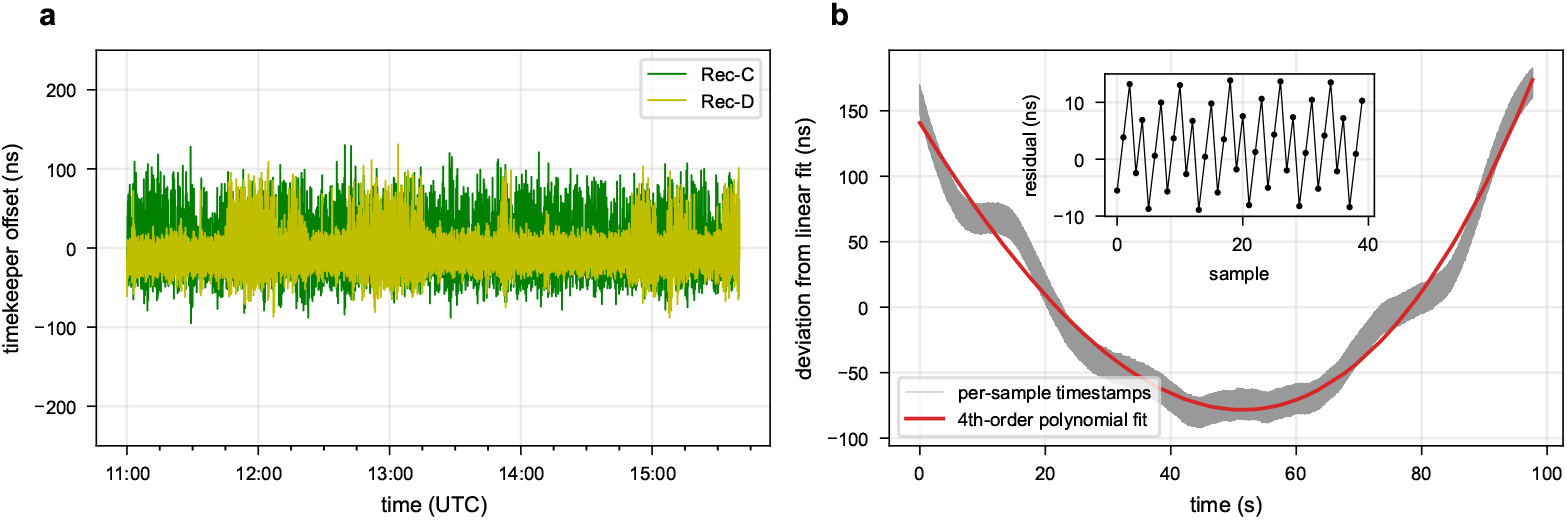
Buoy recorder synchronisation. **a** GPS-disciplined timekeeper offset from two buoy recorders (Rec-C, Rec-D) during one excursion. The timekeeper time stays within ±100 ns of the GPS time. **b** Per-sample timestamp deviation from the nominal 102.4 kHz sampling grid (grey) for 10 × 10^6^ consecutive samples ( 98 s) of Rec-D, after removing a linear drift, with the fourth-order polynomial fit (red). The residual structure reflects higher-order clock accelerations between the GPS clock and the ADC oscillator. Inset: fit residual over 40 consecutive samples, dominated by the 25 ns quantisation of GPS timestamps.

The NI-9250 sigma-delta converters ran on their own quartz oscillator, shared across both modules via the CompactRIO chassis backplane, providing sample-synchronous acquisition across all four channels. The interchannel phase mismatch is specified to 0.035° at 1 kHz, corresponding to 97 ns—the intra-tetrahedron timing precision.

Per-sample timestamps are the basis for the offline resampling pipeline, implemented in the `whalesonic` Python package, which proceeds in three steps. First, a time mapping is established: for each batch of 10^7^ samples (≈98 s), a fourth-order polynomial is fitted to map nominal ADC sample times to UTC, absorbing oscillator drift and the 25 ns quantisation of GPS timestamps (Fig. 4b). Second, a uniform output time grid at 100 kHz is constructed, aligned to UTC second boundaries. Third, the raw signal is interpolated onto this grid using a quartic (*k*=4) univariate spline. Consecutive batches overlap by 2000 samples to ensure continuity at batch boundaries. No additional anti-aliasing filter is applied, because the ADC’s built-in sigma-delta decimation filter limits the signal bandwidth to below the output Nyquist frequency of 50 kHz.

The resulting data are aligned to UTC and directly combinable with other time-referenced data sources, freeing users from clock model and synchronisation concerns. Figure 4 illustrates both levels of the timing chain: the real-time GPS disciplining and the offline resampling to a uniform UTC grid.

#### Data consolidation

Consolidation was performed per excursion in a dedicated Jupyter notebook for reproducibility. First, all GPS tracks from the four buoys, the research vessel, and the speaker were loaded and plotted on a timeline to visualise when each recorder was active. The sound data were then inspected by viewing spectrograms and listening to audio to identify time segments in which all recorders were in the water, stable, and capturing acoustically interesting content. For each selected segment, the resampled audio streams from all available buoys were assembled into a single HDF5 file, together with GPS tracks (resampled to 1 Hz and aligned to UTC) for each buoy and the research vessel, sound velocity profile data, and per-recording metadata. The complete consolidation workflow is implemented in the open-source `whalesonic` Python package.

### Sound emission system

An underwater speaker (LL916, Lubell Labs Inc.) served as the acoustic source for bottom hydrophone synchronisation, localisation, and buoy array validation. The speaker was driven by an amplifier (TPA3116D2, Texas Instruments; 100 W Class-D) connected to a NI-9260 digital-to-analogue converter (National Instruments; 24-bit, 51.2 kHz). As in the buoy recorder system, a GPS time replica running on an FPGA timestamped every output sample with sub-microsecond accuracy. The speaker is a piezoelectric transducer with a usable frequency range of 200 Hz to 23 kHz; beam width narrows with increasing frequency. The amplifier–speaker chain exhibited harmonic distortion, visible as harmonics in the spectrogram during frequency sweeps.

The 21 s test sequence comprised: an opening synchronisation pulse, two white noise segments (5 s each, at different amplitudes), a 5 s linear frequency sweep, four pure tones (1 s each), and a closing synchronisation pulse. The waveform was identical across all emissions, except that the white noise segments were generated independently for each emission. Playback was triggered with the first sample following a UTC second boundary. For each emission, the system recorded the full output wave-form with GPS timestamps for each sample and the geographic track. The arrival of an emission at a receiver was detected by cross-correlating the white noise segment with the received signal and computing the impulse response power (IRP)—the squared envelope of the cross-correlation. The first peak in the IRP corresponds to the direct-path arrival; its timing yields the acoustic propagation delay.

The speaker was suspended from the side of the boat at a depth of 4.65 m below the water surface. The GPS antenna was mounted on the cabin roof, ~1 m offset horizontally from the speaker’s overboard position. The volume setting was kept constant throughout the expedition. A total of 180 emissions were performed for bottom hydrophone synchronisation and localisation. The emission waveforms and GPS coordinates are provided in the dataset.

### Sound velocity profiler

Sound velocity profiles are essential for converting acoustic propagation delays to distances, and their spatial and temporal variability is a significant error source in localisation. Sound velocity was measured using a sound velocity profiler (SWiFT, Valeport Ltd, Totnes, UK). The instrument measures sound velocity via a digital time-of-flight sensor (accuracy ±0.02 m*/*s), along with pressure (accuracy ±0.01 % of full scale) and temperature (accuracy ±0.01 °C), from which it derives conductivity, salinity, and density. An integral GPS receiver records the geographic position and UTC time for each profile.

The profiler was attached to a 200 m cord on a hand reel and lowered from the boat at ~2 m*/*s until it reached the seafloor or the full cord length was deployed, then reeled back on board. A profile was acquired for every buoy deployment and for each bottom hydrophone synchronisation occasion, providing local, contemporaneous sound velocity data. In total, 118 profiles were collected across the study area over the full deployment period (Fig. 5a). Regular profiles above the hydrophone clusters enabled monitoring of temperature and sound velocity trends at the bottom hydrophone sites (Fig. 5b,c).

**Fig. 5.**
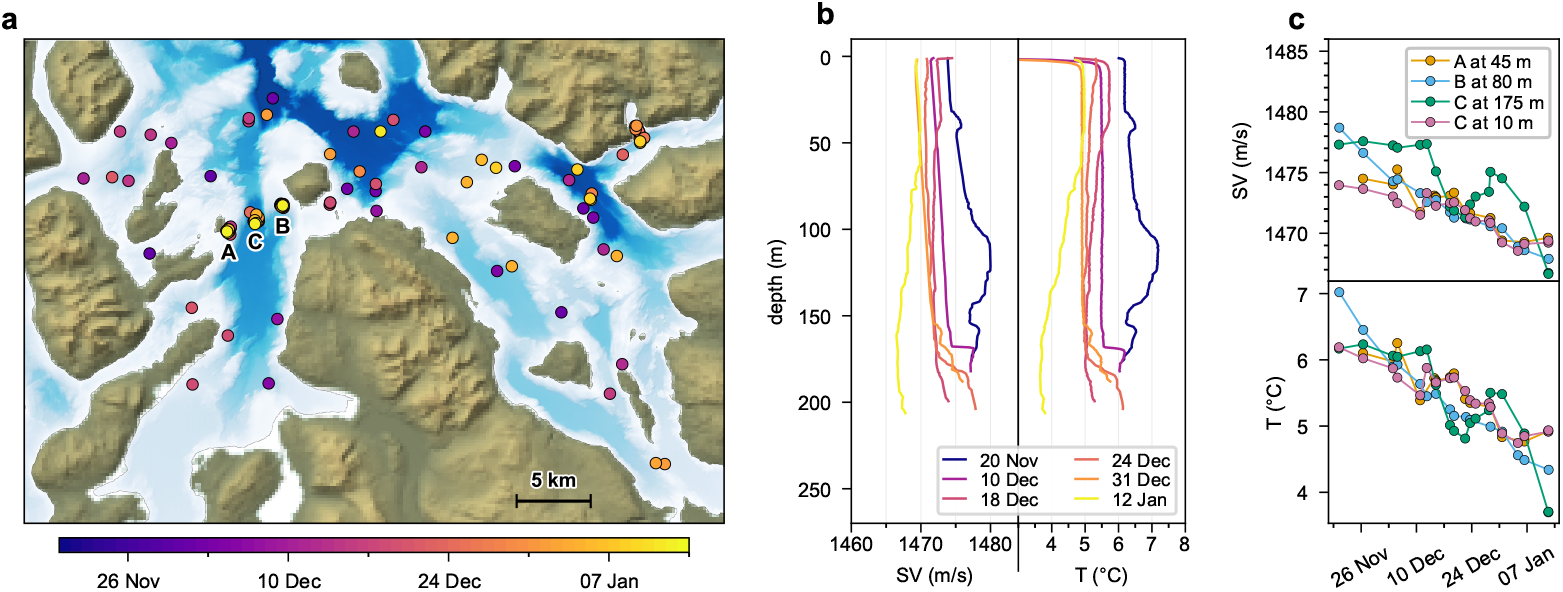
Sound velocity profiles. **a** Locations of 118 sound velocity profiles (dots, coloured by date) across the study area; letters mark the three hydrophone clusters. **b** Six example sound velocity (left) and temperature (right) profiles from station C, evenly spaced across the expedition and coloured by date. **c** Sound velocity (SV) (top) and temperature (bottom) over time at one depth per cluster close to the hydrophones. An additional trace (pink) shows station C at 10 m for comparison with near-surface conditions relevant to the buoy recorders.

The profile data are stored in the dataset without resampling, preserving the instrument’s native sampling resolution. The reported depth values are relative to the water surface at the time of measurement and therefore referenced to the instantaneous tidal water level.

### Bottom hydrophone array

Seven bottom-mounted autonomous recorders provided continuous recording at fixed positions, forming a time-synchronised and acoustically localised array (Fig. 6a). The array was deployed between the islands of Hakstein and Haukøya at the entrance to Reisafjord, a location chosen for its proximity to Skjervøy—enabling visits even in rough weather—and because it monitors a passage where cetacean activity had been observed in previous years. The hydrophones were distributed across two clusters (A1–A3 and B1–B3) and one isolated position (C). Cluster A was on the shallow shelf at depths of 43 m to 66 m, cluster B was deeper at 78 m to 101 m, and station C was in the deep basin at ~288 m. Inter-element spacings were ~200 m within each cluster and ~2 km between adjacent clusters.

**Fig. 6.**
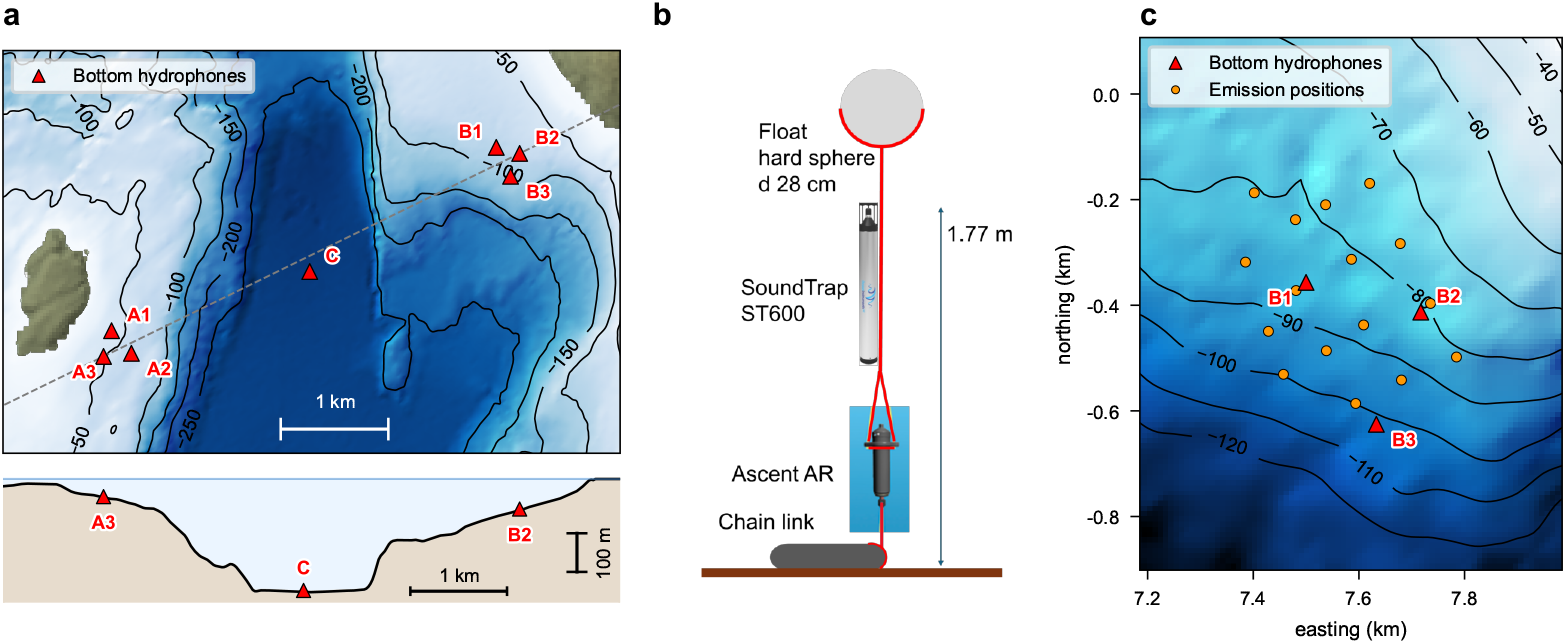
Bottom hydrophone array. **a** Bathymetric map showing the positions of all seven bottom-mounted hydrophones, grouped into cluster A (A1–A3), cluster B (B1–B3), and isolated station C. The dashed line indicates the A3–B2 transect; the cross-section below shows the seafloor profile and hydrophone depths along this line. **b** Schematic of the mooring assembly: steel anchor, acoustic release, recorder, and hard sphere float. **c** Detail map of cluster B showing the 16 sound emission positions (orange circles) used for acoustic localisation.

#### Hardware and mooring assembly

The autonomous recorders (SoundTrap ST600, Ocean Instruments) recorded continuously at a native sample rate of 192 kHz with 16-bit resolution, storing seamless 30 min files to a 512 GB SD card in each unit. The usable acoustic bandwidth extends from 20 Hz to 60 kHz (±3 dB), with factory-calibrated end-to-end sensitivities ranging from −177.3 to −175.5 dB re 1 V*/*µPa across the seven units (Table 3, Table 4).

**Table 3.**
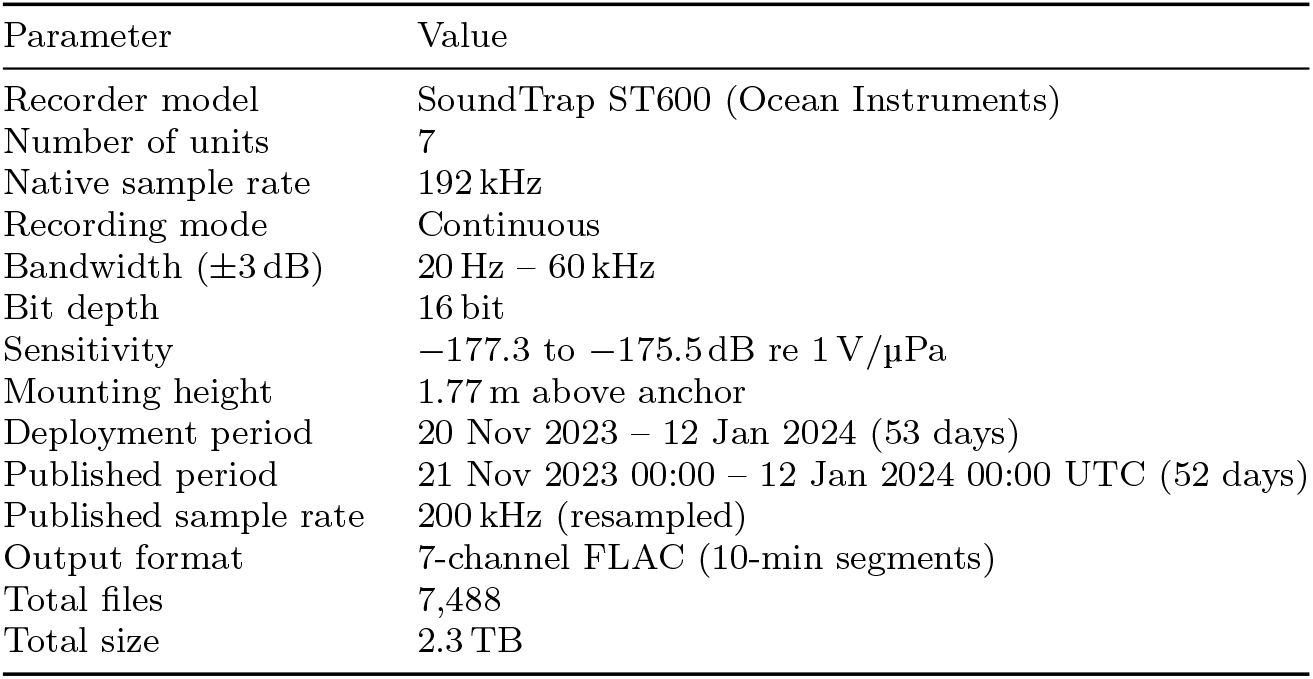
Bottom hydrophone array specifications.

| Parameter | Value |
|---|---|
| Recorder model | SoundTrap ST600 (Ocean Instruments) |
| Number of units | 7 |
| Native sample rate | 192 kHz |
| Recording mode | Continuous |
| Bandwidth (±3 dB) | 20 Hz – 60 kHz |
| Bit depth | 16 bit |
| Sensitivity | −177.3 to −175.5 dB re 1 V/µPa |
| Mounting height | 1.77 m above anchor |
| Deployment period | 20 Nov 2023 – 12 Jan 2024 (53 days) |
| Published period | 21 Nov 2023 00:00 – 12 Jan 2024 00:00 UTC (52 days) |
| Published sample rate | 200 kHz (resampled) |
| Output format | 7-channel FLAC (10-min segments) |
| Total files | 7,488 |
| Total size | 2.3 TB |

**Table 4.** Bottom hydrophone summary. Positions from acoustic localisation in local stereographic coordinates (see Methods). Sensitivity from factory end-to-end calibration. Clock drift rate and acceleration from quadratic polynomial fit to bucket synchronisation and field resynchronisation measurements (*N*); total drift is the accumulated clock error relative to UTC over the 53-day deployment. For B3, eight pre-displacement measurements were excluded from the fit (see B3 mooring displacement). Localisation RMS from least-squares multilateration using 16 sound emissions per hydrophone cluster (Fig. 7**c**).

| ID | Serial | Sens.<br>(dB re 1 V/*μ*Pa) | *x*<br>(m) | *y*<br>(m) | *z*<br>(m) | Loc. RMS<br>(m) (*N* = 16) | Clock drift<br>(ppm) | Accel.<br>(ppb/day) (*N*) | Total drift<br>(s) |
|---|---|---|---|---|---|---|---|---|---|
| A1 | 7922 | −176.0 | 3924 | −2061 | −43 | 1.37 | 2.28 | −0.5 (22) | 10.4 |
| A2 | 7923 | −177.0 | 4109 | −2271 | −66 | 0.77 | 3.16 | +1.7 (22) | 14.4 |
| A3 | 7926 | −176.7 | 3850 | −2300 | −46 | 0.93 | 2.54 | +0.3 (22) | 11.6 |
| C | 7925 | −176.9 | 5766 | −1511 | −289 | 0.56 | 2.65 | +0.3 (24) | 12.1 |
| B1 | 7924 | −175.5 | 7500 | −357 | −84 | 1.39 | 1.68 | +1.1 (19) | 7.7 |
| B2 | 7927 | −177.3 | 7717 | −413 | −78 | 1.41 | 2.23 | −1.5 (19) | 10.2 |
| B3 | 7928 | −176.2 | 7633 | −625 | −101 | 1.55 | 3.06 | +2.2 (11<sup>†</sup>) | 14.1 |

<sup>†</sup>Eight pre-displacement measurements excluded due to mooring displacement (see B3 mooring displacement).

Each recorder was deployed on an individual mooring assembly (Fig. 6b). The mooring consisted of a 15 kg steel anchor, an acoustic release (Ascent AR, Innova Sea), the recorder, and a hard sphere float (28 cm diameter). The hydrophone was mounted 1.77 m above the anchor, with the float at ~3 m to 4 m above the anchor.

#### Synchronisation and localisation principle

The autonomous recorders depended on their internal clock, which drifted by several seconds over the deployment. We synchronised and localised them in three stages. First, the clock offset of each recorder was measured against GPS time immediately before deployment and after recovery, yielding two reference points that defined a linear clock model. Second, using this time model, acoustic emissions from GPS-tracked positions were used to estimate the three-dimensional position of each hydrophone by multilateration. Because the localisation was performed just a few hours before the recovery synchronisation, the time error is negligible. Third, additional acoustic emissions during the deployment—at dates between the two endpoint synchronisations—provided intermediate clock offset measurements that revealed deviations from the linear model and allowed refinement to a higher-order polynomial.

#### Deployment and field operations

All seven recorders were deployed on 20 November 2023 and recovered on 12 January 2024, spanning 53 days. Sound velocity profiles were measured at each cluster site during both deployment and recovery to provide accurate travel-time calculations.

For endpoint clock synchronisations, a bucket of seawater was placed directly on the speaker on deck, the hydrophones were immersed, and the test sequence was played. Cross-correlation of the emitted and recorded signals determined each recorder’s clock offset to within ~±10 µs (equivalent to ±15 mm).

For acoustic localisation, 16 emissions were played by the GPS-linked speaker on a 4 × 4 grid around each cluster on the recovery day, within one to two hours of the post-recovery bucket synchronisation (Fig. 6c).

During the deployment, 15–22 field resynchronisation emissions per cluster were performed opportunistically from the boat. Unlike the bucket synchronisations and recovery-day localisation, which were done in calm conditions, these were not always in ideal weather, and the speaker position could be offset by several metres from the GPS antenna due to boat drift.

#### Data consolidation

The consolidation of the raw data into the published dataset followed the pipeline described below, with each step documented in the `whalesonic` Jupyter notebooks.

##### File integrity

Raw SUD files (SoundTrap proprietary format) were downloaded and FLAC audio extracted. Continuous, uninterrupted sampling is a prerequisite for the synchronisation procedure. We verified that all seven recorders sampled continuously across file boundaries without gaps. The “sampling time” of each recorder is established by taking the timestamp of the first file and counting samples forward. One exception—a ~32-minute gap in recorder A3—was filled with silence to maintain the sample-counting scheme (see A3 SD card anomaly below).

##### Clock synchronisation

The two bucket sync measurements—separated by 53 days—defined a linear clock model for each recorder, mapping sampling time to UTC. Clock drift rates across the seven units ranged from 1.7 to 3.2 ppm, all within the specification of ±5 ppm for the clock (SiT1552 TCXO). Over the 53-day deployment, the accumulated clock offsets ranged from 7.7 to 14.4 s (Fig. 7**a**; Table 4).

**Fig. 7.**
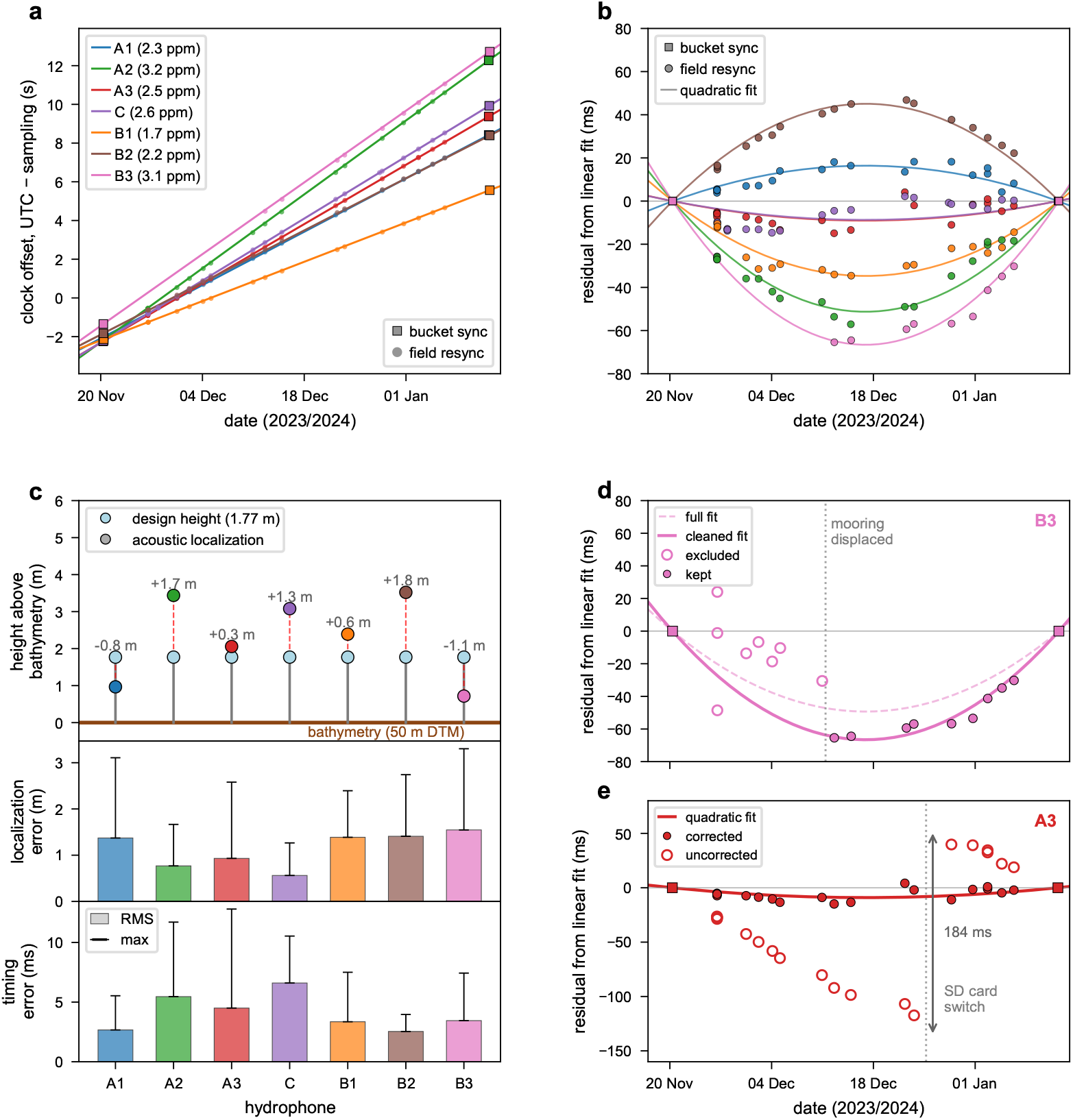
Bottom hydrophone clock synchronisation and localisation. **a** Accumulated clock offset (UTC minus sampling time) over the 53-day deployment, with linear fits through bucket synchronisation points (squares) and field resynchronisation measurements (circles) for all seven hydrophones. **b** Residual after subtracting the linear component, revealing a small quadratic drift corrected by the polynomial model. Markers as in **a. c** Summary of localisation and timing quality across all hydrophones. Top: comparison of acoustically estimated height above the seafloor with the mechanical design height (1.77 m). Middle: localisation residuals (RMS bars, maximum whiskers) from 16-emission multilateration. Bottom: timing residuals from the quadratic clock model. **d** B3 mooring displacement correction. Residual of the B3 clock offset from a linear fit through the two bucket synchronisations. The dashed curve shows the quadratic fit using all measurements; the solid curve shows the cleaned fit after excluding eight pre-displacement resynchronisations (hollow circles). The vertical dotted line marks the displacement on 11 December 2023. **e** A3 SD card switch timing correction. Residual from the corrected linear fit, with the corrected quadratic model (solid curve). Hollow circles show where each resynchronisation measurement would fall without the 184 ms correction; filled circles show the corrected values. The vertical dotted line marks the SD card switch on 25 December 2023.

##### Acoustic localisation

Using the linear clock model, acoustic travel times from the recovery-day emissions (16 per cluster) were extracted by cross-correlation. The linear time model is accurate at this point, just hours before the bucket sync, and travel-time errors are negligible. Combined with sound velocity profiles, least-squares multilateration yielded the three-dimensional position of each hydrophone (Table 4). The localisation residuals range from 0.56 m to 1.55 m RMS (Fig. 7**c**). As a consistency check, the acoustically estimated heights above the bathymetric seafloor range from 0.72 m to 3.52 m, compared with the mechanical mounting height of 1.77 m. The remaining discrepancies are expected, because the 50 m-resolution bathymetry grid does not capture fine-scale bottom topography.

##### Clock model refinement

With the positions now established, the field resynchronisation emissions provided additional clock offset measurements at intermediate dates. The hydrophone positions were assumed known, and the observed travel times yielded clock offsets that revealed a small but measurable drift acceleration (0.3–2.2 ppb*/*day; Table 4), which a linear model cannot capture. Fitting all synchronisation measurements with a second-order polynomial per recorder yielded the final quadratic clock model. However, two anomalies had to be resolved before the final fit could be computed.

#### B3 mooring displacement

On 11 December 2023 10:00, the mooring of hydrophone B3 was displaced ~80 m by fishing operations—confirmed by anchor scraping and rope contact sounds audible in the recording. Because the eight pre-displacement field resynchronisation measurements assumed B3 was at its post-displacement (localised) position, they yielded incorrect clock offsets. Excluding these eight measurements from the clock model fit reduced the B3 timing residual from 19.8 ms to 3.4 ms RMS (Fig. 7**d**), consistent with the other six hydrophones. The localisation and bucket synchronisations are unaffected, as they were performed after the displacement or on deck. The published B3 position is therefore valid from 11 December 2023 onward; before the displacement, the position is unknown within ~80 m.

#### A3 SD card anomaly

Recorder A3 was inadvertently equipped with a 256 GB SD card in its first slot, which filled on 25 December 2023, triggering a switch to the secondary card. This resulted in one truncated file and one absent recording cycle (~30 min gap). The resynchronisation measurements revealed an additional 184 ms timing offset in all post-gap data. The recorder manufacturer confirmed that an SD card switch triggers a reboot, during which the real-time clock is re-read with 1 s resolution, losing sub-second precision. The optimal correction of 184 ms was determined by sweeping candidate values from 0 to 450 ms in 1 ms steps and minimising the RMS residual of the quadratic clock model, reducing it from 41.7 ms to 4.5 ms (Fig. 7**e**). In the published data, channel A3 contains zeros from 2023-12-25 04:58:24 to 05:28:29 UTC.

#### Final clock model and resampling

With both anomalies resolved, all synchronisation measurements were fitted with a second-order polynomial per recorder, yielding the final quadratic clock model that maps sampling time to UTC over the entire deployment. The resulting timing residuals are 2.5–6.6 ms RMS across all seven hydrophones (Fig. 7**b,c**). The quadratic correction was clearly necessary and could be precisely constrained by the resynchronisation data.

Using the quadratic clock model, a regular output time grid at 200 kHz (5 µs sample period) was constructed. The output rate of 200 kHz was chosen over the native 192 kHz for a convenient integer-microsecond sample period that simplifies alignment to the UTC time grid; no spectral content is implied above the native Nyquist frequency of 96 kHz. The raw samples were interpolated onto this uniform grid using quintic interpolation, which provides smooth reconstruction for the slightly higher output sample rate. The DC component was removed by subtracting the mean of each processing chunk.

The published data span 21 November 2023 00:00 to 12 January 2024 00:00 UTC (52 complete days). The deployment and recovery days (20 November and 12 January) are excluded because the recorders were partially on the boat, recording private speech. However, these recordings were used for clock synchronisation and position estimation; the complete procedure can be reproduced from raw data using the provided notebooks. The resampled data were written as seven-channel FLAC files with accompanying metadata.

### Data management

The path from raw instrument data to the published dataset follows a three-tier architecture. The first tier is the *raw data*: the files as they came out of the devices—audio, GPS logs, crew log reports, and photographs—archived together after each excursion in a write-protected repository. These raw data are not published but are retained for full reproducibility.

The second tier is the *consolidation code*: the open-source `whalesonic` Python package and its accompanying Jupyter notebooks. The consolidation process—curation, synchronisation, resampling, calibration, and data fusion—abstracts all instrument-specific technical details and produces measurements in physical units. Every parameter and processing step is documented in the notebooks, which are provided with their saved outputs so that each step can be inspected and reproduced.

The third tier is the *published dataset*: calibrated physical measurements stored in HDF5 and FLAC files with embedded metadata. This separation ensures that users need no knowledge of the acquisition hardware; all instrument-specific complexity is resolved in the consolidation step and can be independently verified through the open-source code.

### Data Records

The dataset is deposited at the ETH Research Collection (DOI: 10.3929/ethz-b-000722140) [56] and comprises buoy array recordings, bottom hydrophone recordings, environmental data, acoustic test signals, and tutorial notebooks (Fig. 8).

**Fig. 8.**
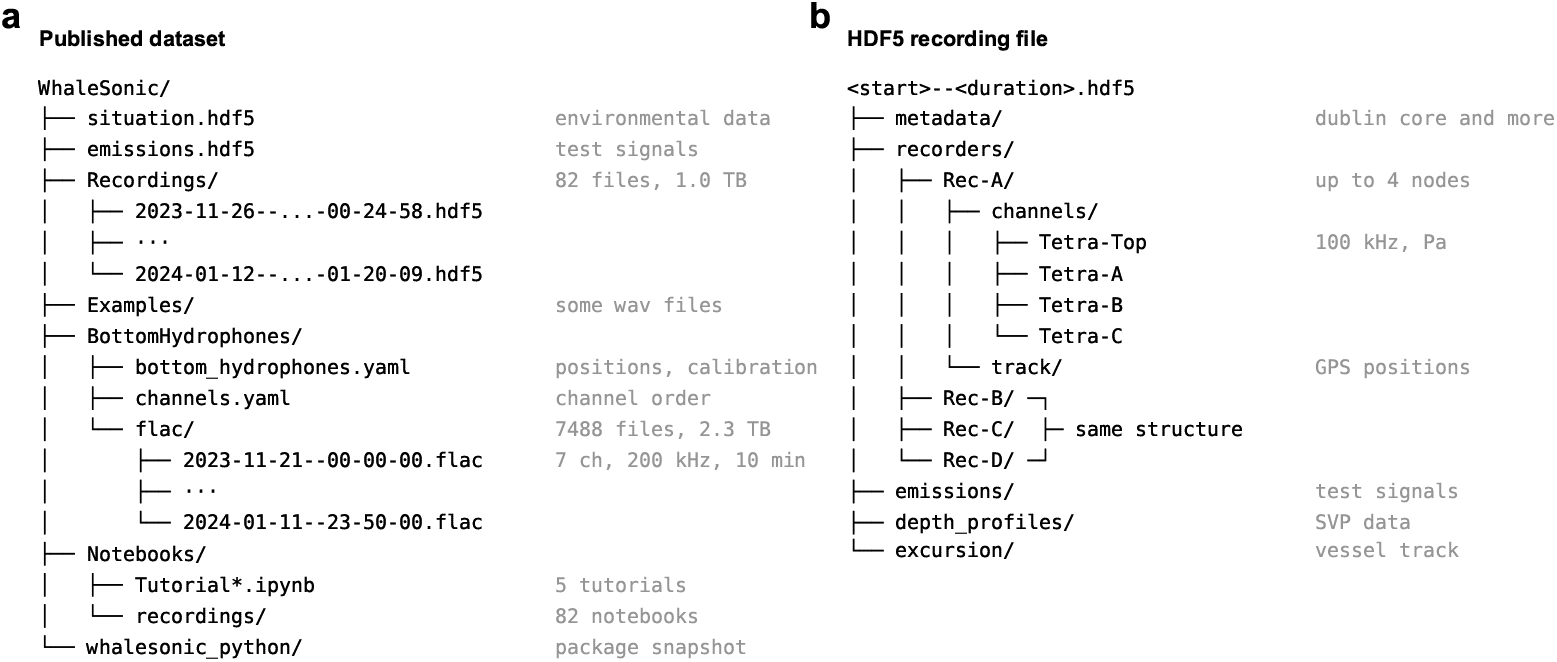
Data organisation. **a** Published dataset directory tree. Buoy array recordings are stored as HDF5 files; bottom hydrophone recordings as 7-channel FLAC files with YAML metadata. Tutorial notebooks, example WAV files for listening, and a snapshot of the whalesonic Python package are included. **b** Internal structure of an HDF5 buoy recording file. Each file contains up to four recorder nodes with four calibrated hydrophone channels, GPS tracks, and optional emission waveforms, depth profiles, and vessel track data.

#### Buoy array recordings

The 82 recordings are stored as individual HDF5 files, named by start time and duration (e.g., `2023-11-26--11-28-02--00-24-58.hdf5`). Each file contains synchronised acoustic data from up to four recorder nodes (Rec-A through Rec-D), with four hydrophone channels per node (Tetra-Top, Tetra-A, Tetra-B, Tetra-C). GPS tracks of each buoy and the research vessel are included, along with sound velocity profile data. If emissions occurred during the recording, the 21 s emission output waveforms are stored in the HDF5 file together with their GPS-referenced timing and position. Attributes at each level of the HDF5 hierarchy make each file self-documenting. All recordings start and end on a UTC second boundary and have a duration that is a whole number of seconds. The 82 HDF5 files total 1.0 TB.

#### Bottom hydrophone recordings

The continuous bottom array data consist of 7,488 seven-channel FLAC files (16-bit, 200 kHz), each covering a 10 min segment. Files are named by start time; the duration is fixed at 10 min and not encoded in the filename. The seven channels are ordered A1, A2, A3, C, B1, B2, B3. The first file is `2023-11-21--00-00-00.flac` and the last is `2024-01-11--23-50-00.flac`, spanning 52 complete days. A YAML metadata file provides per-hydrophone positions, clock synchronisation parameters, and calibration data. The position of hydrophone B3 is valid from 11 December 2023 onward; before this date, the mooring was displaced ~80 m by fishing operations (see Methods). The 7,488 FLAC files total 2.3 TB.

#### Environmental data

The file `situation.hdf5` provides: bathymetry resampled to a 10 m grid in the local stereographic coordinate system; tidal water levels; sound velocity profiles; excursion tracks; and the coordinate system definition enabling conversion between geographic (WGS84) and local coordinates. The `TutorialSituation` notebook documents the contents and usage.

#### Acoustic test signals

The file `emissions.hdf5` contains all speaker emission waveforms and their GPS-referenced positions, enabling acoustic propagation analysis.

#### Tutorial notebooks

Jupyter notebooks with saved outputs are included with the dataset as visual documentation. Tutorial notebooks cover the environmental data (`TutorialSituation`), buoy recordings (`TutorialRecording`), bottom hydrophone data (`TutorialBottomHydrophones`), and data export to WAV or FLAC files (`TutorialExport`). In addition, each of the 82 buoy recordings is accompanied by a per-recording notebook that serves as a visual catalog entry, showing the long-term spectral average, example spectrograms, deployment map, and channel health summary. An `Examples` directory provides eight short stereo WAV clips (≈2 min each, 48 kHz) for direct listening with a standard audio player. A recording overview spreadsheet lists all 82 buoy recordings with deployment context, content summaries, and curated species selections. The `AudioHighlights` notebook documents 46 additional segments selected for their acoustic quality, which can be exported locally from the full dataset. A snapshot of the `whalesonic` Python package is included to ensure reproducibility with the published notebook outputs.

## Technical Validation

### Recording completeness

The buoy recording system produced 82 recordings totalling 78.3 h of multi-channel acoustic data. Each recording can have up to 16 channels (four hydrophones on each of four recorders). Individual channels are classified as *ok* (present and reliable), *broken* (present but exhibiting sporadic distortions or noise due to a bad contact), or *absent* (excluded due to bad quality throughout the recording). Of the 82 recordings, 50 (51.2 h) have all 16 channels fully operational. A further 8 recordings (7.1 h) have all 16 channels present but with one or more channels exhibiting sporadic distortions, bringing the total with 16-channel coverage to 58 recordings (58.3 h). An additional 8 recordings have all four recorder nodes deployed but with some channels absent, for a total of 66 recordings (65.6 h) suitable for long-baseline localisation where just one hydrophone is needed per buoy. The remaining 16 recordings have fewer than four nodes, typically because one or two recorders were undergoing maintenance. Every recording contains at least one complete tetrahedron, suitable for spatial sound field analysis.

Detailed per-recording channel status is available in the per-recording notebooks included with the dataset. Two recordings contain short NaN gaps where a recorder momentarily dropped out: one on 5 December 2023 (Rec-D, 5.1 s) and one on 11 January 2024 (Rec-B, 146 s). All other channels in the affected recordings remain intact.

For the bottom hydrophones, the 7,488 FLAC files span 52 complete days with no gaps. The one exception is channel A3, which contains 30.1 min of silence on 25 December 2023 due to an SD card switch (see Methods).

### Buoy recording system characterisation

Three bench experiments were performed independently of field data to characterise the system noise floor, verify the voltage calibration, and measure the timing offset between two buoys. The first used recorder Rec-B alone; the second and third used the sound emission system’s digital-to-analogue converter (DAC, NI-9260) connected directly to a recorder’s ADC input via a cable, bypassing the speaker and hydrophones.

#### Noise floor

On recorder Rec-B, one ADC input was shorted to measure the converter noise floor, while two channels had their hydrophones placed in a sound-isolated enclosure to measure the combined hydrophone and preamplifier noise (Fig. 9). The NI-9250 ADC exhibited a flat noise floor of 10.3 µV RMS 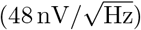, within the manufacturer specification of 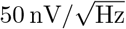, yielding an ADC dynamic range of ~111 dB over the 200 kHz to 41 kHz bandwidth. The hydrophone preamplifier noise measured 40 dB re 1 *µ*Pa^2^/Hz at 1 kHz, 2 dB below the HTI-96-Min specification of 42 dB. Both noise spectra are flat with no spurious peaks across the full bandwidth. The NI-9250 input clips at ±5 V peak. For the −165 dB hydrophones (Rec-B, Rec-C), this corresponds to a clipping level of 179 dB re 1 µPa (peak); for the −170 dB hydrophones (Rec-A, Rec-D), the clipping level is 184 dB re 1 µPa (peak). The system dynamic range is ~92 dB. The noise floor curves in Fig. 9 were measured on Rec-B (−165 dB group); the equivalent curves for the 170 dB group are shifted upward by ~5 dB in acoustic units.

**Fig. 9.**
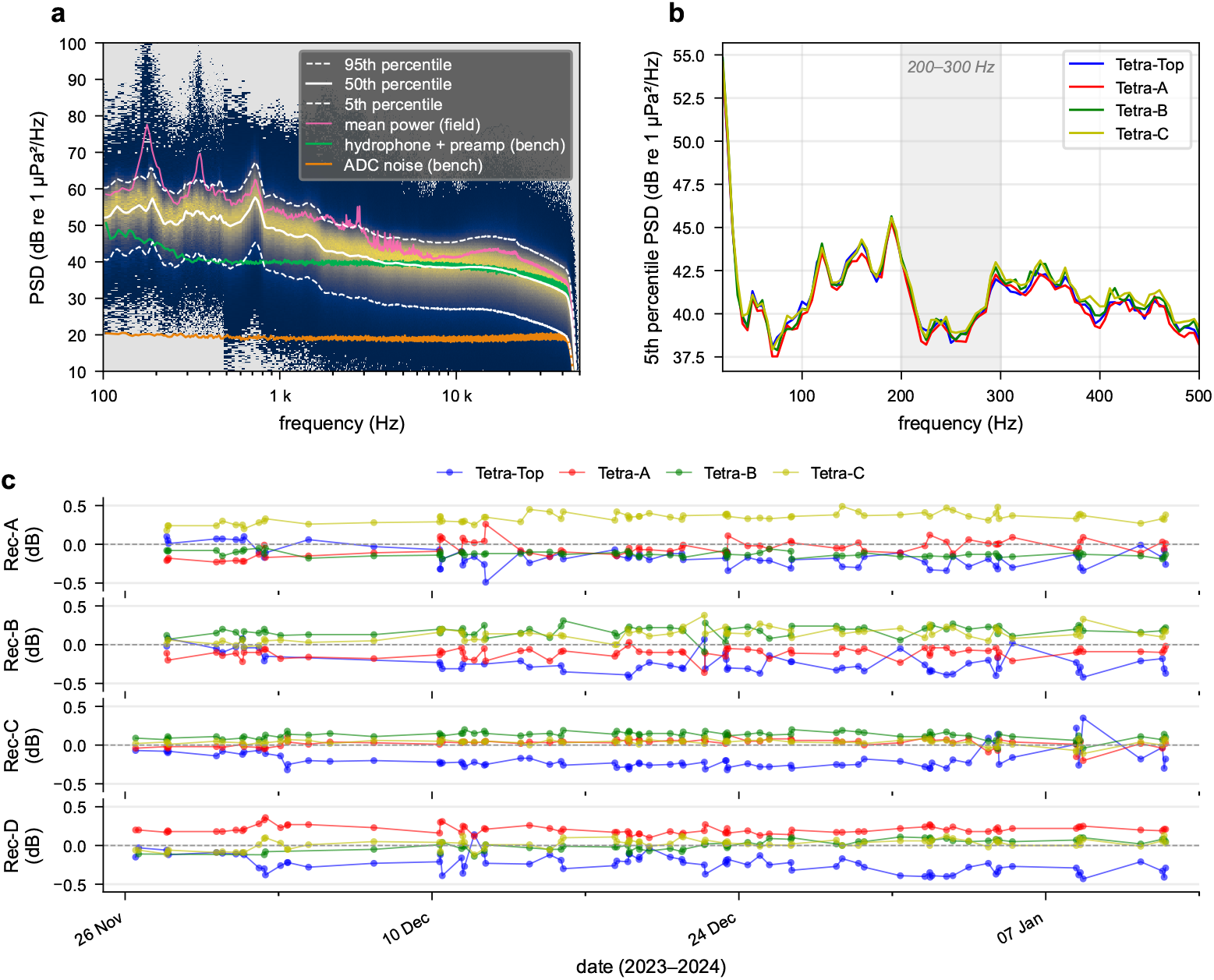
Buoy system characterisation: noise floor and channel gain stability. **a** Spectral probability density function (SPDF) of an example field recording (Rec-B/Tetra-Top, 31 December 2023), where colour indicates the probability of observing a given power spectral density (PSD) at each frequency (dark blue = rare, yellow = frequent). White curves show the 5th, 50th, and 95th percentiles (computed in dB). The bench-test ADC noise floor (shorted input; orange) and combined hydrophone and preamplifier noise (sound-isolated enclosure; green) are overlaid. The Welch mean power spectral density (pink) lies above the 50th percentile because it is a linear-domain average. **b** Low-frequency detail (20–500 Hz) showing the 5th-percentile SPDF curves for all four channels of Rec-B. The shaded band (200–300 Hz) is the frequency range used to extract relative gain offsets. **c** Relative gain offsets over the full expedition for all four recorders. Connected points show each recording; the offsets are small (0.16 dB RMS) and temporally stable (0.07 dB scatter), confirming that channel-to-channel differences are fixed properties of each recording chain.

#### Voltage calibration

The combined DAC/ADC transfer function is flat at 0 dB ±0.1 dB across the 1 kHz to 20 kHz passband, confirming the voltage calibration of both converters.

#### Converter delay

The combined DAC/ADC converter delay was measured at 950 µs, consistent with the 939 µs expected from the NI-9260 and NI-9250 datasheets; the remaining 11 µs is attributed to data transfer to and from the FPGA. This delay was identical across all four buoys and across repeated measurements at different locations and in different years, confirming the accurate and reliable GPS-time synchronisation across independent systems.

The complete bench-test analysis is documented in the `V3 NoiseAndDelay` notebook.

#### Channel gain stability

Because all four hydrophones in a tetrahedron are closely spaced, they receive essentially the same ambient noise field at low frequencies where the acoustic wavelength far exceeds the array dimensions. We exploited this to assess relative channel gains using the spectral probability density function (SPDF). In the 200 Hz to 300 Hz band, the 5th-percentile curve of the SPDF—representing the quietest periods—serves as a gain reference: systematic offsets between channels relative to their geometric mean (zero in dB) reflect the recording chain rather than the acoustic scene.

Applying this analysis to all 82 recordings across the seven-week expedition (documented in the `V4 ChannelGainStability` notebook) yields relative gain offsets that are small and temporally constant. Across all 16 channels, the maximum absolute offset is 0.34 dB, the RMS offset is 0.16 dB, and the mean temporal scatter is 0.07 dB.

### Transmission experiments

To validate the position accuracy of the GPS-tracked buoy array, we compared the acoustically estimated speaker-to-receiver distances with the GPS-measured distances between the speaker and each buoy. During the recovery of the bottom hydrophones on 12 January 2024, we deployed the four buoy recorders near each cluster site and emitted 16 test sequences from a 4 × 4 grid of boat positions (Fig. 10). The resulting 192 transmissions (48 emissions received by four recorders) spanned speaker-to-buoy distances from 20 m to 908 m. The buoy array had an initial spacing of ~200 m but drifted up to about 1 km during the emission series.

**Fig. 10.**
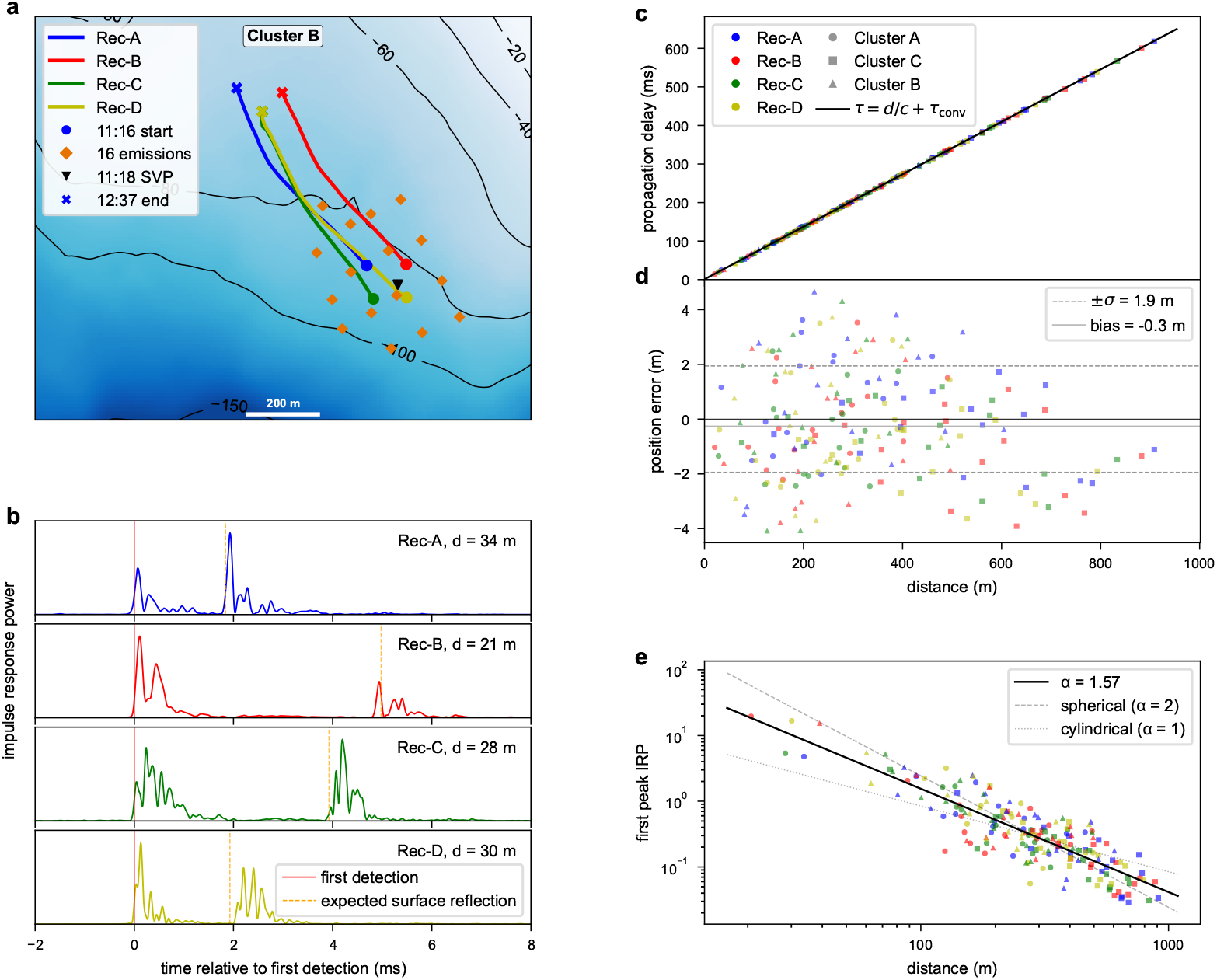
Transmission experiment validation. **a** Overview map of the transmission experiment at cluster B, showing buoy recorder tracks (coloured lines) with start (circle) and end (cross) markers, emission positions (orange diamonds), and sound velocity profile location (SVP, triangle). **b** Close-range impulse response power (IRP) for each recorder’s nearest transmission, showing the first detection (red) and expected surface reflection (orange dashed). **c** Measured propagation delay versus distance for 192 transmissions with the predicted line *τ* = *d/c* + *τ*_conv_. **d** Position error (acoustic minus GPS distance) versus distance, with ±*σ* (dashed) and mean bias (solid grey). **e** First peak IRP versus distance on logarithmic axes with a power-law fit (solid), and reference slopes for spherical (dashed) and cylindrical (dotted) spreading.

#### Positioning accuracy

For each transmission, the propagation delay was measured by detecting the first arrival in the impulse response power (IRP; see Sound emission system). The sound velocity for each emitter–recorder pair was computed from the depth-profiler measurements (accuracy ±0.02 m*/*s; range 1467.6 m*/*s to 1469.3 m*/*s across the three clusters), and the DAC/ADC converter delay of 950 µs was taken from the bench-test measurement (see Converter delay). The difference between the acoustically inferred distance and the GPS-measured distance gives the position error Δ*d* for each transmission. The 192 measurements yield a position error scatter of *σ* = 1.9 m and a mean bias of −0.3 m (RMS 2.0 m; Fig. 10**c, d**). The small negative bias indicates that GPS-based distances are on average slightly longer than acoustic distances. Contributing factors include the ~1 m horizontal offset between the boat’s GPS antenna and the speaker’s overboard position, wind-induced speaker swing of several metres from the hull with a preferred direction (the boat consistently faced into the wind), and horizontal offsets between surface GPS antennas and submerged tetrahedrons due to tidal and wind drift.

The parameter uncertainties—sound velocity (±0.02 m*/*s), converter delay (~10 µs), and GPS timing (*<*1 µs)—contribute less than 0.02 m to the error budget and are negligible compared to the GPS receiver accuracy and antenna-to-transducer offsets.

#### Surface reflections

At close range, the impulse responses clearly resolve the direct arrival and the surface reflection as separate peaks. The predicted reflection delay—computed from the emitter and receiver depths, the horizontal range, and the detected direct-arrival time—matches the observed secondary peak, confirming the geometric consistency of the position and depth data (Fig. 10**b**).

#### Spreading loss and source level

The peak IRP of the first arrival—a measure of the received signal energy at the direct-path arrival time—decays with distance as *r*^−1.57^ (Fig. 10**e**), intermediate between spherical spreading (*r*^−2^) and cylindrical spreading (*r*^−1^), as expected for a shallow-water waveguide where the surface reflection merges with the direct arrival at increasing range. Extrapolating the power-law fit to the 1 m reference distance yields a speaker source level of 184 dB re 1 µPa at 1 m (equivalent speaker sensitivity 200 dB re 1 µPa*/*V at 1 m), averaged over all directions and the 200 Hz to 10 000 Hz analysis bandwidth. The detailed analysis is provided in the `V1 TransmissionExperiments` notebook.

### Species overview

Visual inspection of spectrograms confirmed the presence of four cetacean species in both recording systems (Fig. 11). Each buoy recording file embeds a written content summary based on crew logs and a systematic review of all recordings; a tabular overview with species selections is provided as a spreadsheet (see Data Records). Killer whale calls and echolocation clicks are the most prevalent signals, present in 68 of 82 recordings. Humpback whale vocalisations appear in 50 recordings, sperm whale clicks in 39, and fin whale signals in 31. Engine noise from the research vessel or other boats is audible in most recordings, and anthropogenic sounds from fishing and whale-watching vessels are common, reflecting the high-traffic nature of the study area. A few recordings with little or no biological activity were retained for baseline diversity. The bottom hydrophones, recording continuously, captured the full temporal range including overnight and multi-day sequences not accessible from the buoy system. Fin whale and humpback whale signals are present throughout the 52-day record, typically received on all seven hydrophones. Killer whale sounds appear more frequently in the first half of the deployment and are often detected on only one cluster. Sperm whale clicks are also regularly present.

**Fig. 11.**
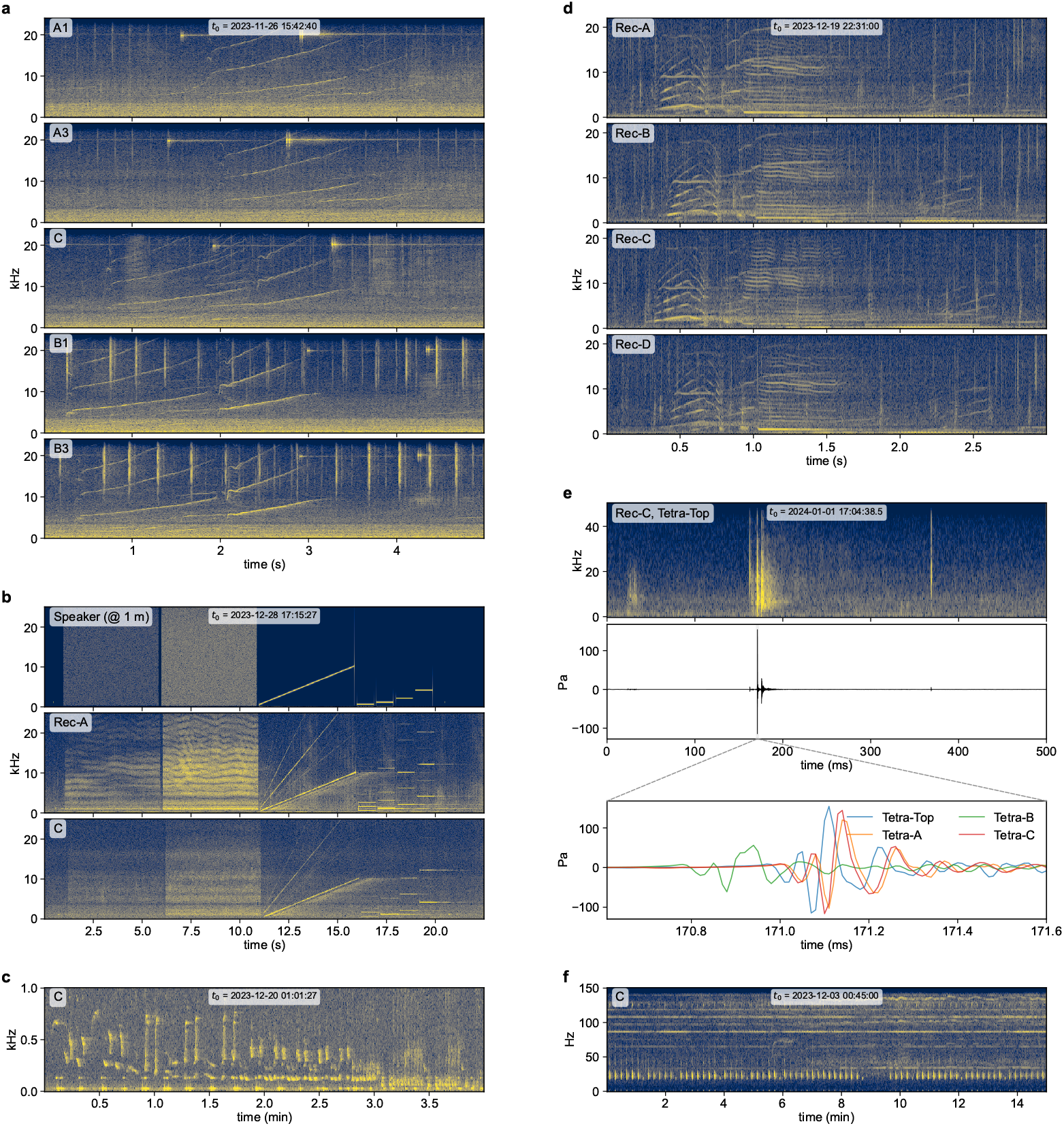
Example recordings from bottom hydrophones, buoy recorders, and the combined system. **a** Killer whale calls recorded simultaneously by five bottom hydrophones, showing time delays across the array. **b** Emission test sequence: speaker output at 1 m (colour scale reduced for clarity) and received signals on buoy recorder Rec-A and bottom hydrophone C, showing propagation delay. **c** Humpback whale song on bottom hydrophone C (signal present on all channels). **d** Killer whale calls recorded simultaneously by four buoy recorders, showing spatial variation across nodes. **e** Sperm whale click on Rec-C tetrahedron: spectrogram (Tetra-Top), full waveform (Tetra-Top), and 1 ms zoom of all four tetrahedron channels showing time-of-arrival differences. Dashed lines indicate the zoom region. **f** Fin whale patterned 20-Hz pulse sequence on bottom hydrophone C (power reduced by 20 dB to fit colour scale). All spectrograms use a 30 dB to 90 dB re 1 *µ*Pa^2^/Hz colour scale unless noted.

### Usage Notes

#### Self-documenting files

The HDF5 files [64] of the WhaleSonic dataset carry their own hierarchical metadata— coordinates, sensitivities, sample rates, units—at every level, making them self-documenting and accessible with any HDF5-compatible tool. Desktop viewers such as HDFView (https://www.hdfgroup.org/download-hdfview/) or the browser-based myHDF5 (https://myhdf5.hdfgroup.org) provide immediate visual inspection without programming; in Python, the files can be read directly with `h5py`. The internal structure of the recording and situation files is documented in the `HDFrecording` and `HDFsituation` notebooks. The bottom hydrophone recordings are distributed as standard FLAC audio files accompanied by a YAML metadata file that provides sensitivity, position, and clock model parameters for each unit. Together, these formats allow users to work with the data using general-purpose libraries and tools, independent of the `whalesonic` package.

#### The whalesonic package

The `whalesonic` Python package provides convenient access to the dataset and facilitates common data handling routines such as extraction, visualisation, and export. Installation and configuration are described in the README of the repository (see Code Availability). The central concept is the `Segment`—a time window of acoustic data, typically a call or a sequence, loaded into memory for analysis. A Segment preserves the hierarchical node-and-channel structure of the recording, including positions and metadata, and provides built-in routines for spectrogram, waveform, and map visualisation. The same interface serves both buoy recordings and bottom hydrophone data, and can be extended to other datasets that provide a compatible extraction routine, making multi-channel analysis algorithms more general and reusable. At its core, each Segment is defined by a `TimeRange` that specifies the time window and sampling grid. The package includes high-quality resampling routines, so users can work at any target sample rate transparently. The repository also contains the consolidation code and notebooks with outputs that document how the published data were produced; these serve reproducibility but are not needed for working with the dataset. Tutorial notebooks (`TutorialRecording`, `TutorialBottomHydrophones`, `TutorialExport`) provide worked examples as entry points.

#### Export and bioacoustics tools

The `whalesonic` package provides export routines at multiple levels: individual segments can be saved as short WAV clips for focused analysis, while complete buoy recordings or bottom hydrophone channels can be exported as collections of files at a user-specified sample rate and channel selection. These exports serve as the bridge to established bioacoustics tools such as Raven Pro [65] for interactive annotation or PAMGuard [42] for automated detection. Standard audio files carry inherent limitations: file size is bounded, waveforms are scaled to 1 full scale, and only limited metadata can be stored. Continuous data must therefore be split into individual files, the waveform rescaled to a reference level that defines the full-scale amplitude (for example, 10 Pa), and users must keep track of the reference level and channel selection. The export workflow is demonstrated in the `TutorialExport` notebook.

#### Timestamps and open annotations

We recommend that annotations be expressed as absolute UTC timestamps rather than as offsets into a particular export file. The files in the published dataset start at integer-second UTC boundaries, so the filename timestamp—which encodes only integer seconds—is exact. Annotation tools that parse the start time from the filename, as Raven Pro does by default, therefore recover correct absolute timestamps. When exporting custom files, aligning them to second boundaries preserves this property. An annotation made on a single-channel WAV export can then be used to access the same moment in the original dataset at full sample rate and across all channels, enabling analyses that go far beyond the original export—for example, localising a stereotyped call that was annotated on just one channel. We encourage users who produce annotations or detections to share them with absolute UTC timestamps for the benefit of the research community; the unified time base of the dataset ensures that such contributions remain interoperable. The same principle applies to the two recording systems themselves: because both are aligned to UTC and referenced to the same local coordinate system, data from the bottom array and the buoy array can be combined directly.

#### DC offsets and low-frequency content

The NI-9250 ADC is AC-coupled with a 3 dB corner frequency of 0.43 Hz. Nevertheless, each channel shows a small DC offset that has not been removed during data consolidation. Surface gravity waves at 0.2 Hz to 2.5 Hz produce pressure fluctuations that can reach 30 Pa RMS depending on sea state and tetrahedron depth—orders of magnitude larger than the acoustic signals of interest. Fin whale 20-Hz pulses can also dominate the low-frequency energy. Users should apply a high-pass filter before exporting to WAV or computing broadband levels: a cutoff of 10 Hz removes the wave pressure and DC offset while preserving all acoustic content; a cutoff of 40 Hz additionally suppresses the fin whale band. The surface wave analysis is documented in the `V5 SurfaceWavePressure` notebook.

## Data Availability

The WhaleSonic dataset is deposited at the ETH Research Collection (DOI: 10.3929/ethz-b-000722140) [56]. All data are publicly available under a Creative Commons Attribution 4.0 International (CC BY 4.0) licence.

## Code Availability

The `whalesonic` Python package is available under the GPL-3.0 licence at https://gitlab.switch.ch/cetacean-communication/expeditions/whalesonic/whalesonic. A snapshot (version 1.1.0) is included with the dataset. The package provides the complete data consolidation pipeline, a segment-based data access API, and tutorial notebooks.

## Author Contributions

**J.R**.: Conceptualisation, Data curation, Formal analysis, Funding acquisition, Investigation, Methodology, Project administration, Resources, Software, Supervision, Validation, Visualisation, Writing – original draft. **A.B**.: Software, Investigation, Writing – review & editing. **J.C**.: Investigation, Project administration, Supervision. **I.D**.: Investigation, Writing – review & editing. **A.E**.: Conceptualisation, Investigation, Project administration, Supervision, Writing – review & editing. **V.F**.: Investigation, Writing – review & editing. **O.F**.: Investigation, Writing – review & editing. **L.G**.: Investigation, Software, Writing – review & editing. **P.H**.: Investigation, Writing – review & editing. **A.H**.: Investigation, Writing – review & editing. **F.K**.: Investigation, Writing – review & editing. **C.L**.: Investigation, Writing – review & editing. **J.G.M**.: Funding acquisition, Project administration, Investigation, Data curation, Writing – review & editing. **C.M**.: Investigation, Writing – review & editing. **F.S.S**.: Investigation, Writing – review & editing. **R.U**.: Investigation, Writing – review & editing. **K.W**.: Funding acquisition, Investigation, Writing – review & editing. **R.H**.: Conceptualisation, Funding acquisition, Resources, Supervision, Writing – review & editing.

## Acknowledgements

We are deeply grateful for the experience of working in this extraordinary Arctic environment and the beauty of the natural world it revealed.

Marten Bril of Whale2Sea provided generous support for the project. Eve Jourdain of the Norwegian Orca Survey provided operative support and invaluable knowledge about the local killer whale population. Zöe Morange and Ove Pedersen joined us for a memorable day recording sperm whales. Carol Haefliger covered the expedition for Swiss television. Krisztina Balotay and Norbi Bus from Orca Channel provided invaluable local information. We thank the whale watching community of Skjervøy for their cooperation and shared adventures, and the local community for their friendliness and help.

We thank the Swiss Embassy in Norway for navigating the Marine Scientific Research application process; the Norwegian Directorate of Fisheries for granting research permission and coordinating with commercial fishing operations; the Norwegian Maritime Authority for approving our safety concept; and the Norwegian Armed Forces for the necessary permits.

This work was supported by ETH Zürich; the NCCR “Evolving Language” Innovation Grant of the Swiss National Science Foundation (Agreement 51NF40 180888); and the Earth Species Project (ESP) AI for Interspecies Communication Challenge Grant. Travel support for J.M. and P.H. was provided by the A.H. Schultz Foundation. O.F. was supported by Carlsberg Center for Active Sensing with Sound (CF24-2176, PI Peter Teglberg Madsen).

## Ethics Declaration

Passive acoustic recordings do not require ethics approval under Norwegian and European animal research legislation (regulation of 18 June 2015 No. 761, § 2f), as confirmed by the Norwegian Food Safety Authority. Research was conducted under permit from the Norwegian Directorate of Fisheries.

